# Gene function and expression profiling in yeast spores, killifish diapause embryos, and their post-dormant offspring cells

**DOI:** 10.64898/2026.05.08.723705

**Authors:** Shaimaa Hassan, Maria Rodriguez-López, StJohn Townsend, Büşra Köksal, Sezin Akkus, Alessandro Ori, Alessandro Cellerino, Markus Ralser, Jürg Bähler

## Abstract

Dormancy is a reversible cellular state characterised by suspended proliferation and increased stress resilience, enabling long-term viability under adverse conditions. Although dormant cells are critical for the life cycle of diverse organisms, from microbes to humans, they are understudied compared to proliferating cells. We present a comparative investigation of dormant cells in two divergent species: spores of fission yeast and diapause embryos of turquoise killifish. A genome-wide screen for genes affecting the lifespan and heat-shock resilience of spores uncovered a trade-off between longevity and heat resistance, and considerable differences in the genetic basis for lifespan between spores and chronologically aging yeast cells. RNA-seq and mass-spectrometry analyses revealed substantial transcriptomic and proteomic changes in spores and diapause embryos, with ribosomal proteins induced as transcripts but repressed as proteins. Transcriptomic regulation of biological processes, but less so of specific genes, is broadly conserved across yeast spores, killifish diapause, and human dormant cancer cells, including the induction of autophagy– and translation-related processes and the repression of cell cycle-related processes. Spores and diapause embryos modulate their transcriptomes and proteomes in response to heat stress and prolonged time. These RNA and protein expression changes are uncoupled and differ from aging-related expression signatures in yeast cells and adult fish. Cells derived from older or stressed spores retain phenotypic differences for several cell divisions, reflected in altered expression signatures, lifespan and stress resilience. Similarly, diapause duration and heat exposure are associated with long-term expression signatures in post-diapause embryos before hatching. This study highlights core biological processes and principles that are remarkably conserved in distinct types of dormant cells.

## Introduction

Dormancy is a controlled, reversible cellular state of arrested proliferation and reduced metabolic activity across diverse organisms. From unicellular microbes to plants and vertebrates, the ability to delay the life cycle by entering a dormant state is a widespread survival strategy in response to adverse environmental conditions. For example, bacteria and yeast cells respond to nutrient limitation by suspending growth and division as quiescent cells or as spores, which feature increased stress resistance and long-term survival (Rittershaus et al. 2013; de Virgilio 2012; Errington 2003). Dormancy is also an adaptive strategy for many animals (Hand et al. 2016). Examples include nematodes entering a dauer stage in response to poor conditions and the various phases of diapause in insects, fish and mammals to endure predictable periods of harsh environments in different life stages (Fielenbach and Antebi 2008; Gottlieb and Ruvkun 1994; Mead 1981; Ptak et al. 2012; Renfree and Fenelon 2017; MacRae 2010). Dormant cells are also critical for human health and aging. For example, the alternation between quiescent and proliferating cell states plays a pivotal role in stem-cell function, tissue renewal, and immune responses (Oh et al. 2014; Li and Clevers 2010; Marescal and Cheeseman 2020; Valcourt et al. 2012; Elkin et al. 2025). Moreover, long-lived dormant cancer cells can evade therapies targeting proliferating cancer cells and later re-enter the cell cycle, driving relapse (Ebinger et al. 2016; Phan and Croucher 2020; Jiang et al. 2025).

Although dormant cells are widespread in nature, they are far less studied than proliferating cells, and fundamental questions about their biology remain unanswered. For example, to what extent do dormant stages in diverse organisms and contexts reflect conserved biological processes and what factors are critical for the regulation and maintenance of dormancy? There are no molecular markers for dormant cells, reflecting our limited understanding of this state. Here, we analyse dormant stages of two evolutionarily divergent model organisms, the fission yeast *Schizosaccharomyces pombe*, a single-celled fungus, and the turquoise killifish *Nothobranchius furzeri,* a vertebrate. These two species provide complementary, broad perspectives on the genetic basis and cellular regulation during dormancy.

Fission yeast cells will enter a quiescent state in the absence of nitrogen, characterised by a reversible arrest of cell proliferation, increased stress resistance, and reprogramming of gene expression and energy metabolism from a growth mode to a maintenance mode (de Virgilio 2012; Marguerat et al. 2012; Yanagida 2009; López-Maury et al. 2008; Roche et al. 2017; Valcourt et al. 2012; Sideri et al. 2015; Braun et al. 2026). If yeast mating partners are present, nitrogen limitation triggers meiotic differentiation and gene-expression reprogramming (Mata et al. 2002; 2007), culminating in the formation of dormant spores that feature extraordinary stress resilience and longevity (Ohtsuka et al. 2022; Tsuyuzaki et al. 2021, 2020; Sakai et al. 2025). The revival of budding yeast from 200-year-old beer in a shipwreck highlights the astonishing capacity of dormant cells for long-term survival (de Virgilio 2012).

The turquoise killifish has emerged as a compelling vertebrate model for studying aging and associated diseases (Poeschla and Valenzano 2020; Brunet 2020; Platzer and Englert 2016; Cellerino et al. 2016). This annual killifish inhabits seasonal ponds in Southeast Africa that persist only during the rainy season. Remarkably, the fish survives the long dry season in embryonic diapause, a dormant state of arrested development (Harel 2022; Dolfi et al. 2019; Furness et al. 2015; Reichwald et al. 2015). Annual killifish embryos can enter diapause at three distinct developmental stages; these diapauses are optional and of variable length, even within a single clutch, likely reflecting bet-hedging strategies in response to unpredictable environments (Furness et al. 2015; Wourms 1972; Polačik et al. 2014). Diapause II occurs at mid-somitogenesis, when most organs are formed, and is the most prominent dormant state in *N. furzeri* (Platzer and Englert 2016; Hu et al. 2020). Different aspects of gene regulation during *N. furzeri* diapause II have been investigated (Reichwald et al. 2015; Hu et al. 2020; Singh et al. 2024).

Here, we present a cross-species comparative approach to investigate conserved survival strategies associated with dormant cells. We report a large-scale deletion-mutant screen using yeast spores to identify genes and cellular processes required for cell survival and stress resistance during dormancy. We then analyse genome regulation in *S. pombe* spores and *N. furzeri* diapause embryos at both the transcriptome and proteome levels. We identify commonalities in regulated biological processes in different types of dormant states, including published transcriptome data from dormant cancer cells and quiescent yeast cells (Ebinger et al. 2016; Marguerat et al. 2012). We also examine how dormant cells respond at the transcriptomic and proteomic levels to heat stress and prolonged periods in dormancy. Notably, stress and dormancy duration are ‘remembered’ in the active cells after dormancy exit, reflected by lasting gene-expression signatures and cellular traits. These results highlight key regulated processes, post-transcriptional principles, and core genes shared across diverse dormant states. This integrated study from distantly related model systems contributes to a better understanding of the genetic and regulatory basis of dormancy, with implications for both basic biology and biomedicine.

## Results and Discussion

### Genes affecting the longevity and stress resilience of yeast spores

To identify key genes involved in the lifespan and stress resistance of yeast spores, we conducted a large-scale genetic screen using Barcode sequencing (Bar-seq) of *S. pombe* gene-deletion libraries tagged with mutant-specific barcodes (D.-U. Kim et al. 2010; Han et al. 2010). We have previously decoded the barcodes of 3206 mutants in the library (Romila et al. 2021). Using a colony arraying robot, we mixed corresponding haploid colonies from an auxotroph and a prototroph deletion library on malt-extract medium to trigger mating and sporulation, resulting in haploid spores that each contained a defined deletion mutant (Figure 1A). The spores were pooled and aged, with samples collected from 2 weeks to 6 months, as well as after exposure to a 30-minute heat shock at 55°C (Figure 1A). To avoid DNA contamination from dead cells (Romila et al. 2021), the spore samples were germinated in rich medium, and cells were grown to the stationary phase before sequencing (Figure 1A). Two independent biological repeats were carried out for this screen (Pools A and B).

**Figure 1:**
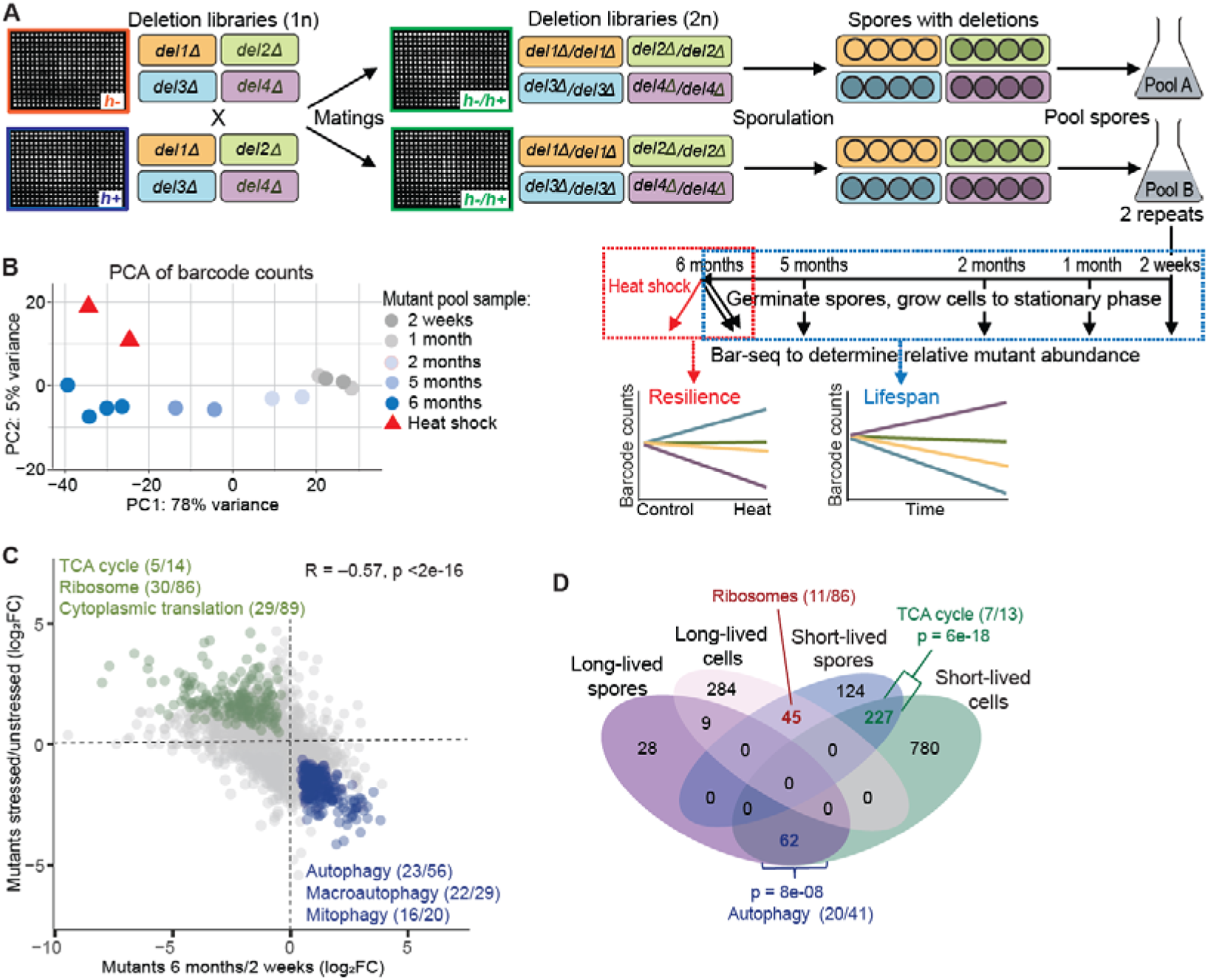
Genetic screen for mutants affecting spore lifespan and stress resilience in *S. pombe*. **(A)** Experimental design for the parallel functional profiling of barcode-tagged deletion mutants using Bar-seq to identify genes required for spore lifespan or heat-stress resilience. Haploid deletion libraries were mated and sporulated; the resulting spores, each containing a specific gene deletion, were pooled, aged for 6 months in water at 25°C, and then an aliquot was heat-shocked at 55°C for 30 minutes. Aliquots were germinated, grown and sequenced at different times, as indicated, to quantify the relative abundance of the barcode-tagged mutant strains. At 6 months, unstressed spores were germinated in 2 technical replicates. The screen was performed in 2 independent biological repeats (Pools A and B). **(B)** Principal Component Analysis (PCA) of barcode counts (upper-quartile-normalised log_2_CPM [counts per million]) among the different repeated pooled spore samples. Dot colors represent the timepoints as indicated, and red triangles represent the heat-shocked samples. The variances represented in the principal components 1 and 2 are indicated on the axes. **(C)** Scatterplot comparing the log_2_ fold-changes (FC) in mutant barcodes in heat-stressed relative to unstressed spore pools and in old (6-month) relative to young (2-week) spore pools. Each point represents a deletion mutant. The Pearson correlation coefficient between spore stress resilience and longevity phenotypes is shown at top right, along with its significance. Green dots: mutants being significantly stress resistant and short-lived; blue dots: mutants being significantly stress sensitive and long-lived (FC >1.5; FDR < 0.05), with other mutants in grey. Enriched GO and KEGG terms are provided for stress-resistant but short-lived mutants (green) and stress-sensitive but long-lived mutants (blue), along with the gene counts for those terms present in the dataset. **(D)** Overlaps of lifespan mutants in stationary-phase cells (Romila et al. 2021) and in spores (this study). For overlaps that are higher than expected by chance, the p-values are indicated (Fisher’s exact test), along with enriched GO terms and gene counts for those terms present in the dataset. Only mutants that were detected in both screens are included in the comparison.

**Figure S1:**
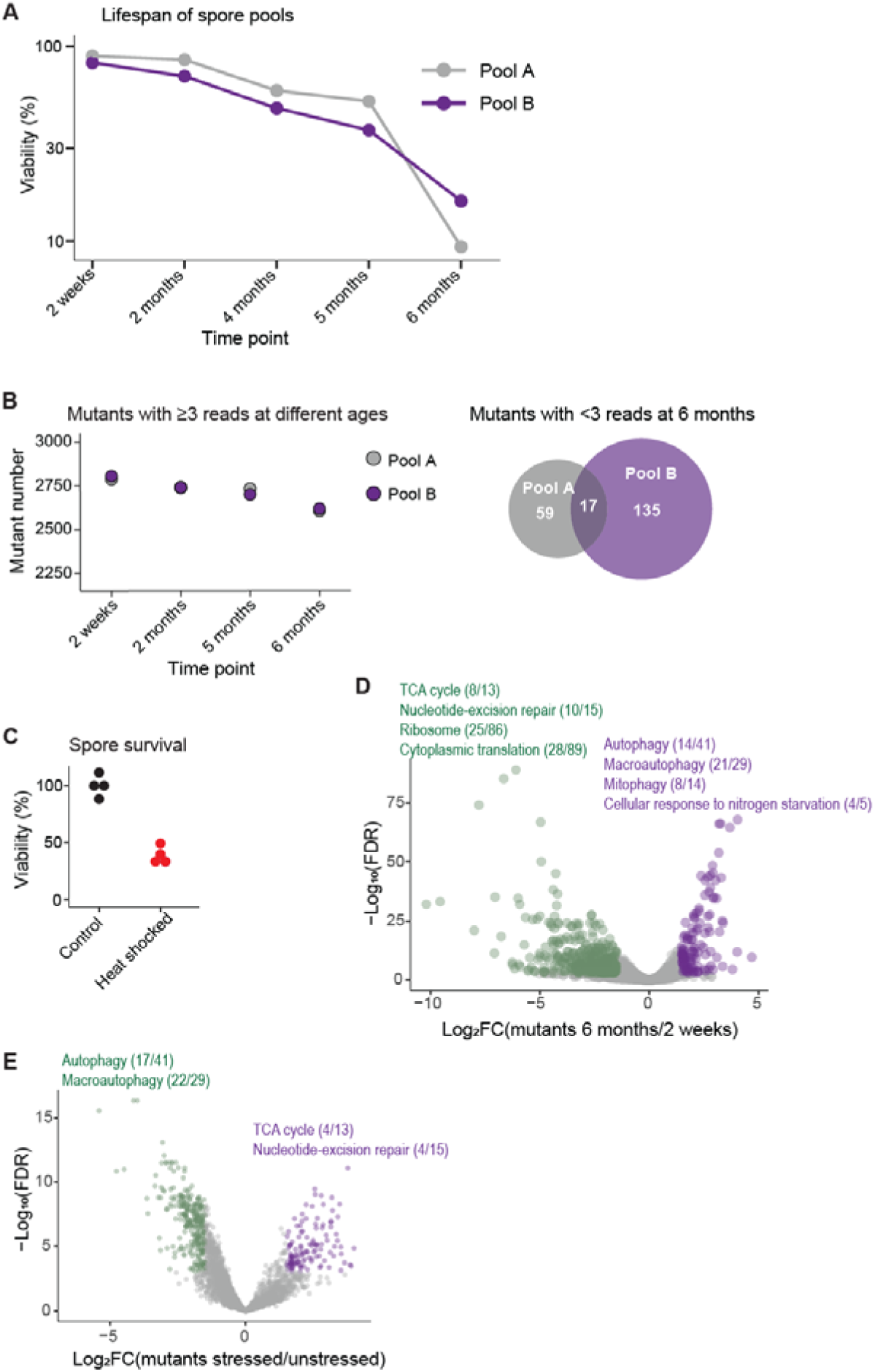
Genetic screen for mutants affecting spore lifespan and stress resilience. (**A**) Viability of pooled spores as a function of time. Pools A and B are the two independent spore pools from the Bar-seq screen (Figure 1A). Mean spore viability was determined at each time by isolating 100 spores using a tetrad dissection microscope, plating them on YES, and counting the resulting cell colonies. **(B)** Left: Number of mutants detected at ≥3 reads/million in Pools A and B at the different time points indicated (of 3,206 mutants in the library). Right: Number of mutants detected at <3 reads/million in Pools A and B after 6 months in any of the 2 technical repeats; these mutants were discarded from the analysis. **(C)** Viability of 2-week-old wild-type spores maintained at 25° (Control) or subjected to a 30-minute heat stress at 55°C (Heat shocked). Each dot represents 1 of 4 independent repeats. **(D)** Volcano plot displaying changes in mutant (barcode) distributions, with normalisation done to the upper quartile. The mutants that significantly decreased or increased >3-fold at 6 months relative to 2 weeks are indicated in green and purple, respectively. Enriched GO terms and KEGG pathways for induced and repressed genes are shown, along with the gene counts for those terms present in the dataset. **(E)** Volcano plot displaying changes in mutant (barcode) distributions. The mutants that significantly decreased or increased >3-fold after heat stress relative to the unstressed condition are indicated in green and purple, respectively. Enriched GO terms and KEGG pathways for induced and repressed genes are indicated, along with the gene counts for those terms present in the dataset.

At 6 months, the average viability of the spores had decreased to ∼10-20% in the two pools (Figure S1A). Of the 3,206 mutants in the library, we initially detected 2,806 and 2,827 mutants (87-88%) with ≥3 sequence reads per million in Pools A and B, respectively. After 6 months, there was only a slight decrease, with 2,730 and 2,675 mutants (83-85%) still detected (Figure S1B). The 55°C heat shock killed ∼60% of wild-type spores (Figure S1C), providing a robust dynamic range. Following the heat shock at 6 months, 2,687 and 2,593 mutants were detected with ≥3 sequence reads per million in Pools A and B, respectively. A Principal Component Analysis (PCA) of the barcode counts, reflecting each mutant’s proportion in the samples, showed variation as a function of time and, to a lesser extent, in response to heat (Figure 1B). PC1 separated the samples by age, with the 5– and 6-month samples being distinct from samples collected earlier. The stressed sample was similar to the 6-month unstressed control sample on PC1 but distinct on PC2. Although the total number of mutants detected remained relatively stable (Figure S1B), these results indicate that the relative mutant composition changed substantially in older samples and that heat induced additional changes distinct from those caused by aging.

To identify mutants whose proportions change with age and/or heat stress, we used edgeR (Robinson et al. 2010; Romila et al. 2021). We treated time as a categorical variable and included a pool factor to account for differences in barcode abundance across the 2 pools, thereby mitigating for library composition effects. A negative binomial generalised linear model with likelihood ratio testing was used to detect significant changes in relative barcode abundance (Romila et al. 2021). Mutants were classified as long– or short-lived by comparing their distribution at 6 months relative to 2 weeks, and as stress-resistant or – sensitive by comparing their distribution at 6 months before and after heat stress, applying thresholds for fold-change >3 and FDR <0.05.

Using this approach, we identified 99 long-lived and 416 short-lived spore mutants. The long-lived mutants were enriched for autophagy-related functions (Figure S1D; Dataset S1). Although autophagy mutants are known to exhibit mating defects in *S. pombe* (Sun et al. 2013; Mukaiyama et al. 2009), the malt-extract medium apparently provided sufficient nitrogen for these mutants to form spores. The 10 longest lived mutants involved 3 autophagy-related proteins (Atg101, Atg8, Ctl1), 3 proteins with mitochondrial functions (Cox18, Mtg1, Nnr2), 2 metabolic proteins (Acb1, Hem14), a chromatin protein (Nhp6), and the trehalase Ntp1. Ntp1 degrades trehalose, required for effective spore germination (Eleutherio et al. 2015; Inoue and Shimoda 1981); thus, the longevity of *ntp1* deletion mutants suggests that increased trehalose is beneficial for dormant spores. The short-lived mutants, on the other hand, were enriched for processes such as the mitochondrial tricarboxylic acid (TCA) cycle, nucleotide excision repair, and ribosomes (Figure S1D; Dataset S1). The shortest lived mutants involved 2 proteins with known roles in spore formation and maturation (Mfr1, Neo1) (Blanco et al. 2001; Ucisik-Akkaya et al. 2014), 2 proteins required for spore fitness in a recent sexual reproduction screen (Plb1, Alg9) (Billmyre et al. 2022), a protein functioning in ribosome biogenesis (Rok1), 3 mitochondrial proteins (Sdh1, Pma2, Emi5), a chromatin protein (Mms19), as well as 2 transcription factors, the CCAAT-binding factor complex component Php3, required for efficient sporulation (Ucisik-Akkaya et al. 2014; Protacio et al. 2022), and Rsv2, which induces stress-related genes during spore formation (Mata et al. 2007).

Regarding the heat shock, we identified 116 resistant and 258 sensitive mutants. The resistant mutants were enriched for TCA cycle and DNA repair functions (Figure S1E; Dataset S1). The most stress-resistant mutants involved mitochondrial proteins (Emi5, Ilv1, Oms1) and DNA repair proteins (Rad13, Rhp14, Rhp26, Rhp41). The stress-sensitive mutants were enriched for autophagy functions (Figure S1E; Dataset S1). The most sensitive mutants involved proteins functioning in lipid and tRNA metabolism (The4, Dus1) (Finet et al. 2022) and in vesicle-mediated transport and detoxification (Erv46, Fmd2) (Astuti et al. 2016), a histone deacetylase complex protein (Png3) (Zilio et al. 2014), and 4 autophagy proteins (Atg5, Atg8, Atg13, Atg1801).

The Bar-seq screens for lifespan and stress resilience yielded similar functional enrichments, albeit in reverse directions (Figure S1D,E). Therefore, we examined the global relationship between mutant fitness and age or stress. Notably, these phenotypes were inversely correlated in pooled assays following germination and regrowth, with long-lived mutants tending to be stress-sensitive, while short-lived mutants tending to be stress-resistant (Figure 1C). TCA cycle and ribosomal proteins typically increased longevity but decreased stress resistance, while autophagy proteins decreased longevity but increased stress resistance (Figure 1C). This surprising finding suggests potential trade-offs between processes that promote spore longevity and those that promote stress resilience under acute heat shock. Energy metabolism, protein translation, and autophagy are critical, interconnected processes with complex and context-dependent roles in maintaining cellular homeostasis and aging (López-Otín et al. 2023). A recent study reported detrimental effects of autophagy in quiescent worm cells, caused by lysosomal damage (Murley et al. 2025). Our screen detected mutants for 87 of the 115 ‘priority unstudied genes’, which are conserved from fission yeast to humans, as annotated in PomBase (September 2024; Wood et al. 2019), and 6 of these mutants were short-lived, 3 were heat-resistant, and 19 were heat-sensitive. These findings suggest that many of these conserved yet unknown genes function in dormant spores, a state that is understudied.

We compared the mutants affecting spore lifespan with those affecting the chronological lifespan of stationary-phase cells, identified in a previous Bar-seq screen (Romila et al. 2021). The short-lived mutants in spores and cells showed a significant overlap and were enriched in TCA cycle proteins (Figure 1D). While there was limited overlap between long-lived mutants in spores and cells, a significant overlap was observed between long-lived mutants in spores and short-lived mutants in cells, with the corresponding genes being enriched in autophagy functions (Figure 1D). Another substantial overlap was observed between short-lived spore mutants and long-lived cell mutants, with enriched ribosomal proteins (Figure 1D). Thus, mitochondrial respiration is important for the longevity of both spores and stationary-phase cells, while ribosome– and autophagy-related functions have opposite effects on the lifespans of spores and cells. The lifespan effects observed in stationary-phase cells, in which autophagy promotes longevity and ribosomes suppress it, better align with those observed for aging in other organisms (López-Otín et al. 2023). This finding suggests that the genetic basis and biological processes underlying longevity in long-term dormant cells differ markedly from those in aging cells and organisms.

### Transcriptome and proteome changes in yeast spores

The regulation of transcriptomes and proteomes in different cellular states offers complementary information to genetic screens (Yeger-Lotem et al. 2009; Malecki et al. 2016). To further characterize dormant *S. pombe* spores, we analyzed gene expression of spores relative to vegetative cells at both transcript and protein levels. To this end, homothallic cells (*h^90^*) were mated and sporulated, and the resulting spores were used to prepare samples for RNA-seq and mass-spectrometry analyses, as well as to germinate proliferating *h^90^* cells as a control (Figure 2A). Three independent biological repeats were carried out, with transcriptome and proteome measurements performed on the same samples to maximise comparability between datasets. Overall, we detected 3722 mRNAs (73.6%) and 2645 proteins (52.4%) of the 5057 annotated protein-coding genes in *S. pombe*, and 1023 (13.4%) of the 7627 annotated long non-coding RNAs in at least 3 of 6 samples. Large proportions of these molecules were differentially expressed in spores relative to proliferating cells: 2555 mRNAs (68.6%), 1083 proteins (40.9%), and 706 long non-coding RNAs (69.0%) (Figure S2A; Dataset S2). The relative changes in mRNA levels showed a moderate correlation with the corresponding changes in protein levels, with 334 genes induced and 536 repressed at both transcript and protein levels (Figure 2B). For the 115 priority unstudied genes (Wood et al. 2019), we detected 84 mRNAs, of which 61 (72.6%) were differentially expressed in spores, and 48 proteins, of which 21 (43.7%) were differentially expressed (Dataset S2).

**Figure 2:**
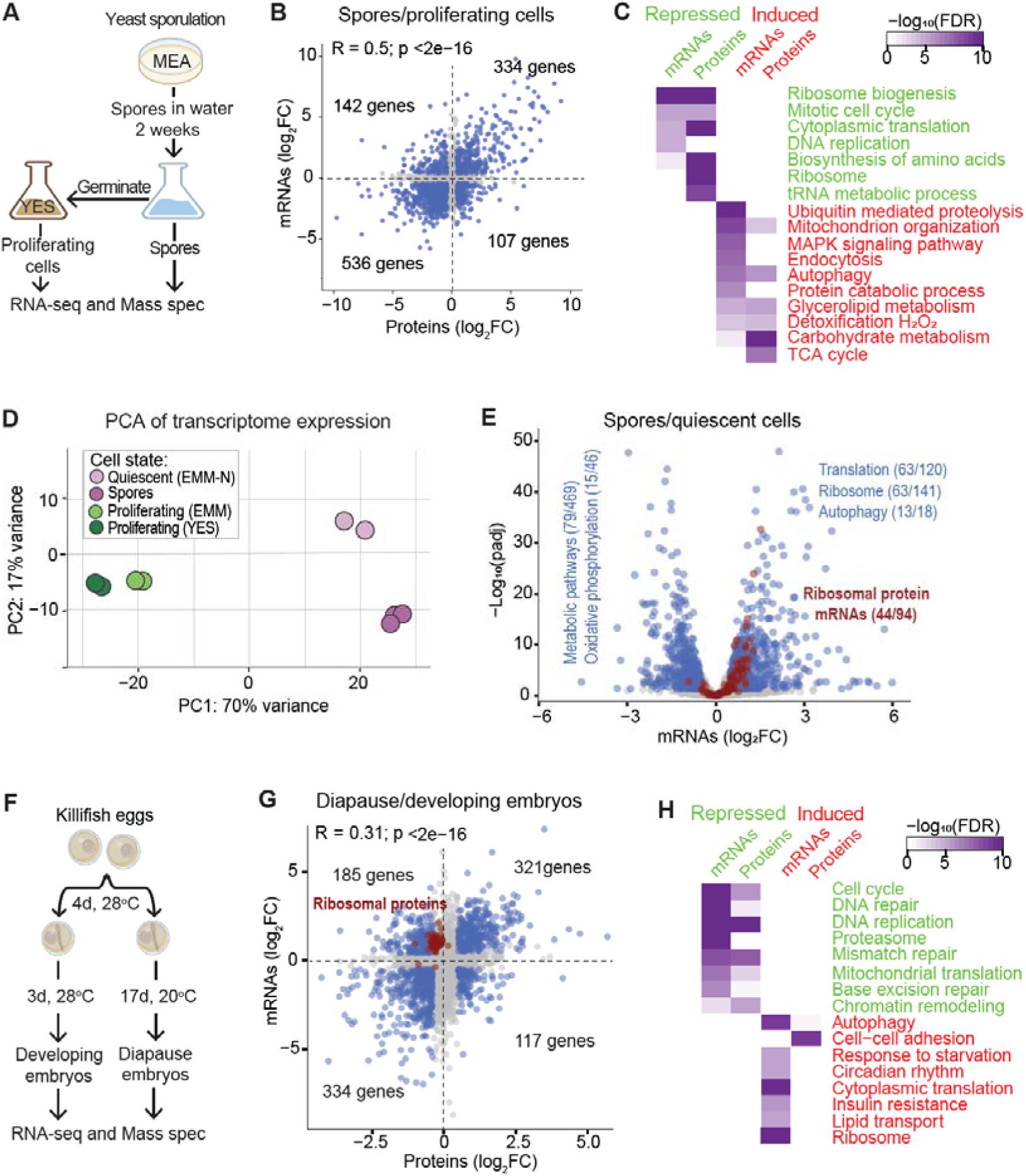
Transcriptome and proteome remodelling in yeast spores and killifish diapause embryos. (**A**) Experimental design for analyzing *S. pombe* spores and proliferating control cells. A homozygous strain was precultured in Yeast Extract Supplements (YES) liquid medium and then in Malt Extract Agar (MEA) plates as indicated. Three independent biological replicates were performed and processed for RNA-seq and mass spectrometry from the same samples. (**B**) Scatter plot comparing the log_2_ fold-changes (FC) in mRNA and protein levels in spores relative to those in proliferating cells. The Pearson correlation coefficient for mRNA and protein changes is shown at the top, along with its significance. The number of genes in each quadrant is indicated. Genes differentially expressed (FC >1.5×; FDR <0.05) for both mRNAs and proteins are highlighted in blue, with other genes in grey. (**C**) Enriched GO Biological Processes and KEGG pathways associated with repressed (green) and induced (red) mRNAs and proteins as indicated. The colour shade corresponds to the false discovery rate (FDR; top right). Enrichments were calculated with g:Profiler2 (Raudvere et al. 2019). (**D**) Principal Component Analysis (PCA) of transcriptomic signatures from spores (this study), quiescent cells (Marguerat et al. 2012), and their proliferating cell controls. Dot colours represent the conditions as indicated. The variances represented in the principal components 1 and 2 are indicated on the axes. The gene expression values used for the PCA are normalized read counts derived from DESeq2. (**E**) Volcano plot showing mRNA expression in spores relative to quiescent cells. Blue dots: mRNAs that are significantly lower (497 genes) or higher (1189 genes) expressed in spores. Enriched GO and KEGG terms for induced and repressed genes are indicated, along with the proportions of genes with those terms. Red dots: 94 mRNAs encoding ribosomal proteins, 44 of which are significantly induced. Significance thresholds for differential expression: FC >1.5×; FDR <0.05. (**F**) Experimental design for analyzing *N. fuzeri* diapause embryos (GRZ strain). Embryos were incubated at 28°C for 4 days post-fertilisation, followed by incubation at 20°C for 14 days to obtain diapause embryos or for an extra 3 days at 28°C to obtain actively developing embryos at a similar stage to the diapause embryos. Three independent biological repeats, collected from different breeding times, were performed and processed for RNA-seq and mass spectrometry from the same samples. (**G**) Scatter plot comparing the log_2_ fold-changes (FC) in mRNA and protein levels in diapause relative to those in developing embryos. The Pearson correlation coefficient for mRNA and protein changes is shown at the top, along with its significance. The number of genes in each quadrant is indicated. Genes differentially expressed (FC >1.5×; FDR <0.05) for both mRNAs and proteins are highlighted in blue, with other genes in grey. All 64 genes encoding ribosomal proteins are highlighted in red. (**H**) Enriched GO Biological Processes and KEGG pathways associated with the human orthologs of the repressed (green) and induced (red) killifish mRNAs and proteins as indicated. The colour shade represents the FDR (top right). Enrichments were determined using g:Profiler2.

**Figure S2:**
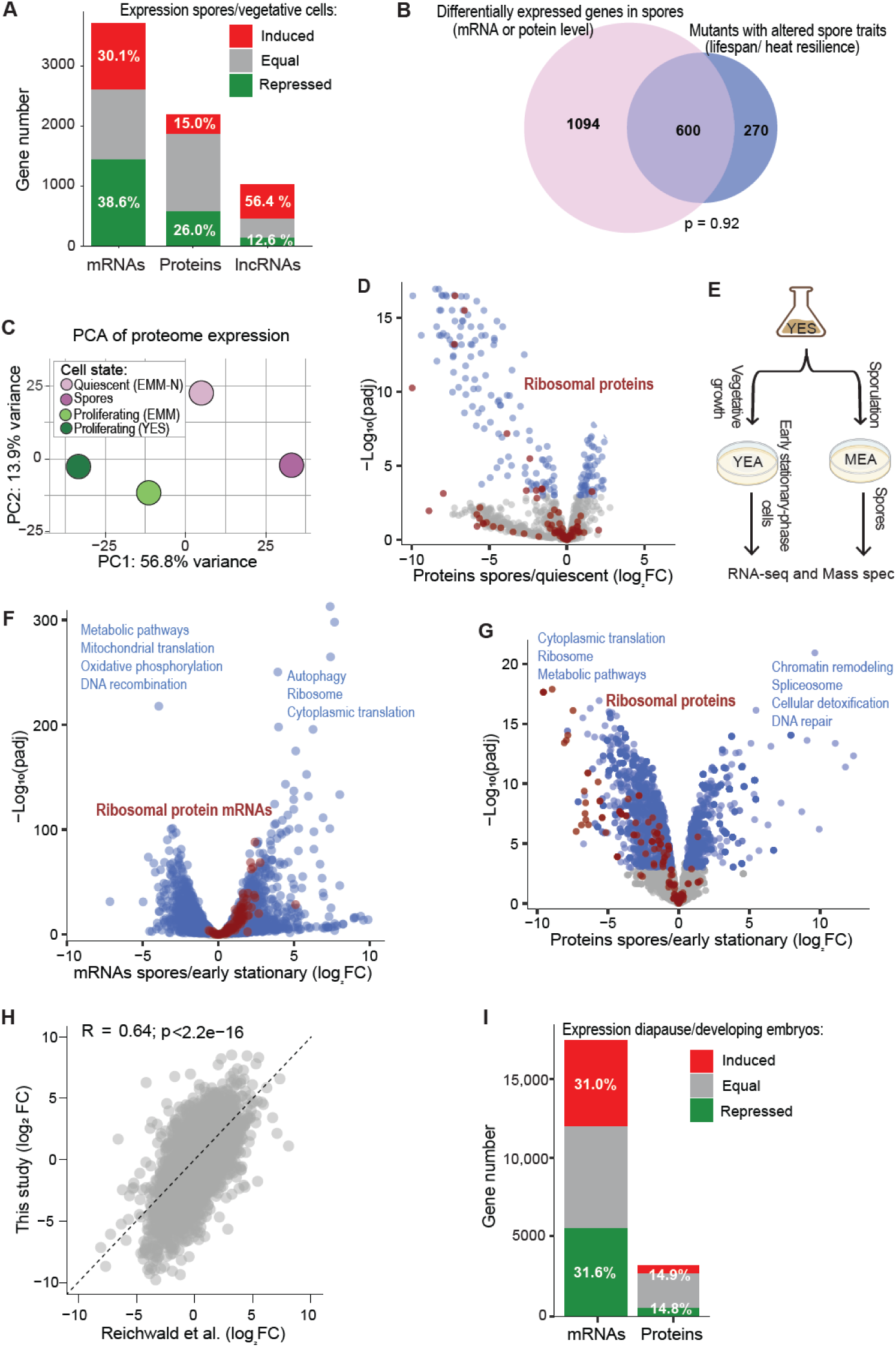
Transcriptome and proteome remodelling in yeast spores and killifish diapause embryos. (**A**) Bar plots showing the numbers and proportions of mRNAs, proteins, and long non-coding RNAs (lncRNAs) being differentially expressed in spores relative to proliferating cells, colour-coded for genes that are induced (red), repressed (green), or non-differentially expressed (grey). Significance thresholds for differential expression: FC >1.5x; FDR <0.05. **(B)** Overlap between mutants with altered spore heat resistance, spore longevity, and differentially expressed genes at RNA or protein levels in spores relative to exponentially growing cells. Only the 2396 genes detected in both Bar-seq and expression data (RNA or proteome) are included. The probability of enrichment in the overlap is included (hypergeometric distribution), with 615 genes expected to overlap by chance. **(C)** Principal Component Analysis (PCA) of proteomic signatures from spores, quiescent cells, and their proliferating control cells. Each dot colour shows the condition as indicated. The variances represented in the principal components are indicated on the axes. The gene expression values used for the PCA are normalized read counts derived from DESeq2. **(D)** Volcano plot showing protein expression in spores relative to quiescent cells. Blue dots: proteins that are significantly lower or higher expressed in spores. Red dots: ribosomal proteins. Significance thresholds for differential expression: FC >1.5x; FDR <0.05. **(E)** Overview of experimental design for analyzing *S. pombe* spores compared to early stationary-phase cells. A homozygous strain was cultured in Yeast Extract Supplements (YES) liquid medium, Malt Extract Agar plates (MEA) and YES Agar (YEA) plates as indicated. Three independent biological repeats were performed and processed for RNA-seq and mass spectrometry from the same samples. **(F,G)** Volcano plot showing mRNA (F) and protein (G) expression in spores relative to stationary-phase cells. Blue dots: genes that are significantly lower or higher expressed in spores. Enriched GO and KEGG terms for induced and repressed genes are indicated, along with the proportions of genes with those terms. Red dots: ribosomal protein genes. Significance thresholds for differential expression: FC >1.5x; FDR <0.05. **(H)** Scatter plot of the log_2_ fold-change (FC) of mRNA levels in diapause embryos in our data compared to those of Reichwald et al. (2015). The Pearson correlation coefficient for the mRNA changes is shown at the top, along with its significance. **(I)** Bar graph representing the numbers and proportions of mRNAs and proteins being differentially expressed in diapause relative to developing embryos, colour-coded for genes that are induced (red), repressed (green), or non-differentially expressed (grey). Significance thresholds for differential expression: FC >1.5x; FDR <0.05.

We looked for functional enrichments of cellular processes among the differentially expressed genes (Figure 2C). DNA replication, cell cycle, amino acid synthesis, ribosome biogenesis, and protein translation were repressed at the mRNA and/or protein level, while autophagy, proteolysis, MAPK signalling, and mitochondrial functions, including the TCA cycle, were induced (Figure 2C). This analysis highlights that the transcriptomes and proteomes are substantially remodelled in spores, reflecting a relative downregulation of anabolic processes functioning in cell growth and an upregulation of catabolic processes functioning in cell maintenance. Notably, the overlap between genes differentially expressed in spores and those that affect spore lifespan or stress resilience was not higher than expected by chance, even though 600 genes were in common (Figure S2B). This finding is consistent with studies showing that genes regulated in a given cellular state often do not overlap with those functionally required for that state (Yeger-Lotem et al. 2009; Yanai et al. 2005; Malecki et al. 2016). Nevertheless, protein translation, ribosomes, the TCA cycle, and autophagy were strongly enriched processes in both the differentially expressed genes in spores (Figure 2C) and the mutants with altered spore lifespan and stress resilience (Figure 1C). Thus, this comparison reveals key cellular processes that are regulated and functionally crucial in *S. pombe* spores.

We then compared the transcriptomic and proteomic signatures in spores with those in quiescent *S. pombe* cells grown in the absence of nitrogen (Marguerat et al. 2012). A PCA of the transcriptomic signatures revealed that the largest variance between these datasets is driven by differences between proliferating control cells and dormant cells (quiescent cells and spores), while the second-largest variance is driven by differences between quiescent cells and spores (Figure 2D). This result indicates that the transcriptomic signatures of spores and quiescent cells, although distinct, are more similar to each other than those of proliferating cells. The 497 mRNAs expressed at lower levels in spores than in quiescent cells were enriched for metabolic and mitochondrial functions, while the 1189 mRNAs more highly expressed in spores were enriched for functions related to autophagy and cytoplasmic translation, including 44 mRNAs encoding ribosomal proteins (Figure 2E; Dataset S2). The proteome data are sparser but show similar PCA patterns across the cellular states (Figure S2C), with 121 proteins showing lower expression in spores than in quiescent cells and 104 proteins showing higher expression in spores. The ribosomal proteins tended to be expressed at lower levels in spores than in quiescent cells (Figure S2D; Dataset S2), a trend opposite to the corresponding mRNAs (Figure 2E).

We also compared the spore transcriptome with that of vegetative cells in early stationary phase, grown overnight on solid media (Figure S2E,F). The mRNAs showing higher expression in spores than in stationary-phase cells were enriched for functions related to autophagy and cytoplasmic translation, including ribosomal proteins (Figure S2F; Dataset S2). In contrast, the proteins showing lower expression in spores were enriched for ribosomal proteins and other proteins involved in cytoplasmic translation (Figure S2G; Dataset S2). These results are similar to the comparison between spores and quiescent cells (Figure 2E; Figure S2C). We conclude that the transcriptome and proteome signatures of *S. pombe* non-dividing cells and dormant spores exhibit both substantial similarities and clear differences. Most notably, ribosomal proteins and other translational factors show higher expression at the mRNA level but lower expression at the protein level in spores than in quiescent or stationary-phase cells.

### Transcriptome and proteome changes in killifish diapause embryos

To uncover conserved aspects of dormant states between yeast and vertebrates, we analyzed the transcript and protein signatures of *N. furzeri* diapause II embryos, henceforth called diapause embryos, relative to actively developing embryos of the same developmental stage (Figure 2F). Previous studies have conducted transcriptomic analyses of *N. furzeri* diapause embryos (Reichwald et al. 2015; Hu et al. 2020), but no proteomic data have been reported. As for yeast, we extracted RNAs and proteins from identical samples to maximize comparability between transcriptomic and proteomic data. Three independent biological repeats were performed. As the diapause and developing embryos needed to be maintained at different temperatures (Figure 2F), some of the expression differences might reflect temperature rather than the diapause state. To validate our RNA-seq data, we compared them to published data (Reichwald et al. 2015). The relative changes in mRNA levels showed a good correlation between the two datasets, although they tended to be larger in our data (Figure S2H). Overall, we measured 17,595 out of 26,141 annotated mRNAs (67.3%) and 3,243 proteins (12.4%). Large proportions of the measured molecules were differentially expressed in diapause, including 11,018 mRNAs (62.6%) and 963 proteins (29.7%) (Figure S2I; Dataset S2). The relative changes in mRNA levels showed a moderate correlation with the corresponding changes in protein levels, with 321 genes induced and 334 repressed at both levels (Figure 2G). We detected 94 mRNAs and 25 proteins for *N. furzeri* orthologs of priority unstudied *S. pombe* genes. Among these genes, 65 mRNAs (69.1%) and 5 proteins (20%) were differentially expressed in diapause (Dataset S2), again suggesting that many of these unknown proteins may function in dormant cell states.

Notably, 67 genes encoding cytoplasmic ribosomal proteins were induced at the mRNA level but repressed at the protein level in diapause embryos (Figure 2G). An unexpected upregulation of ribosomal protein mRNAs in diapause has also been observed previously (Hu et al. 2020; Reichwald et al. 2015). Here, we show that this induction occurs only at the mRNA level, while the corresponding proteins are repressed. This finding is similar to the relatively higher expression of ribosomal-protein mRNAs, but not proteins, in *S. pombe* spores compared to non-dividing cells (Figure 2E; Figure S2D,F,G). While a decreased need for ribosomal proteins is expected in dormant cells, the relatively high levels of mRNAs for ribosomal proteins could facilitate a rapid increase in protein synthesis upon exit from dormancy when conditions permit. Indeed, the exit from diapause is characterised by a rapid resumption of cell proliferation (Dolfi et al. 2019). Ribosomal mRNAs in dormant cells may be stored and stabilized in a translationally inactive state, e.g. in stress granules (Marcelo et al. 2021) or by selective sequestration in nuclear speckles (Rossi et al. 2026). Notably, in budding and fission yeasts, the splicing of ribosomal protein genes is repressed in quiescent cells (Braun et al. 2026). In mammals, increased levels of mRNAs encoding ribosomal proteins are consistently associated with longevity (Tyshkovskiy et al. 2023). Moreover, an uncoupling in ribosomal-protein regulation at the mRNA and protein level has been reported in the context of aging (Kelmer Sacramento et al. 2020; Janssens et al. 2015). In aging killifish brains, this mRNA-protein decoupling of ribosomal proteins has recently been linked to increased ribosome pausing, thereby stabilizing the mRNAs for ribosomal proteins while decreasing the translation of the corresponding proteins (Di Fraia et al. 2025).

We looked for functional enrichments among the differentially expressed genes (Figure 2H). Cellular processes related to the cell cycle, DNA repair, chromatin remodelling, and mitochondrial translation were repressed at the mRNA and/or protein level, while processes related to autophagy, cytoplasmic translation, starvation response, circadian rhythm, cell adhesion, and lipid transport were induced (Figure 2H). Circadian rhythm genes are also induced in other diapausing animals (A. W. Thompson and Ortí 2016). A recent study has revealed that lipid metabolism plays a unique and central role in *N. furzeri* diapause embryos, which exhibit a distinct lipid profile (Singh et al. 2024). We conclude that, as in yeast spores, the transcriptomes and proteomes are substantially remodelled in killifish diapause embryos. Notably, both spores and diapause embryos show increased expression of mRNAs encoding ribosomal proteins, while the corresponding proteins are repressed, suggesting that these proteins are controlled at the level of translation and/or protein turnover. The conservation of decoupled ribosomal mRNA and protein levels in dormant cells of two evolutionarily distant organisms points to the functional importance of this regulatory pattern.

### Common processes are regulated by gene expression in yeast spores, killifish diapause embryos and human dormant cancer cells

We explored the conservation of genome regulation across distinct dormant cell types in 3 organisms by comparing our data in yeast spores and killifish diapause embryos with published RNA-seq data from dormant human cancer cells (Ebinger et al. 2016). This study sampled patient-derived long-term dormant cells from acute lymphoblastic leukaemia xenografts compared to proliferating cells from the same xenografts (Ebinger et al. 2016). We compared the 2239 human genes with orthologs in the other two organisms, and whose mRNAs were detected in all 3 datasets, applying many-to-many ortholog relationships and showing separate data for paralogs. For yeast and killifish, we also compared our proteome data, which included 1103 orthologous proteins in both species.

The relative expression changes of orthologous mRNAs showed poor, non-significant correlations across the 3 species (Figure 3A). For orthologous proteins, the relative expression changes were significantly, albeit weakly, correlated (Figure 3B). Given the similarities in enriched biological processes among the differentially expressed genes in yeast spores and killifish diapause embryos (Figure 2C and H), we also examined correlations at the process level by averaging the log_2_ fold-changes across all mRNAs or proteins belonging to the same GO Biological Processes. Notably, at this process level, we observed much stronger, significant correlations for all 3 pairwise combinations of yeast, killifish, and human mRNA (Figure 3A) and for yeast and killifish protein data (Figure 3B). This analysis indicates that while the regulation of specific genes within processes often diverges across organisms, the overall regulation of biological processes is much more conserved.

**Figure 3:**
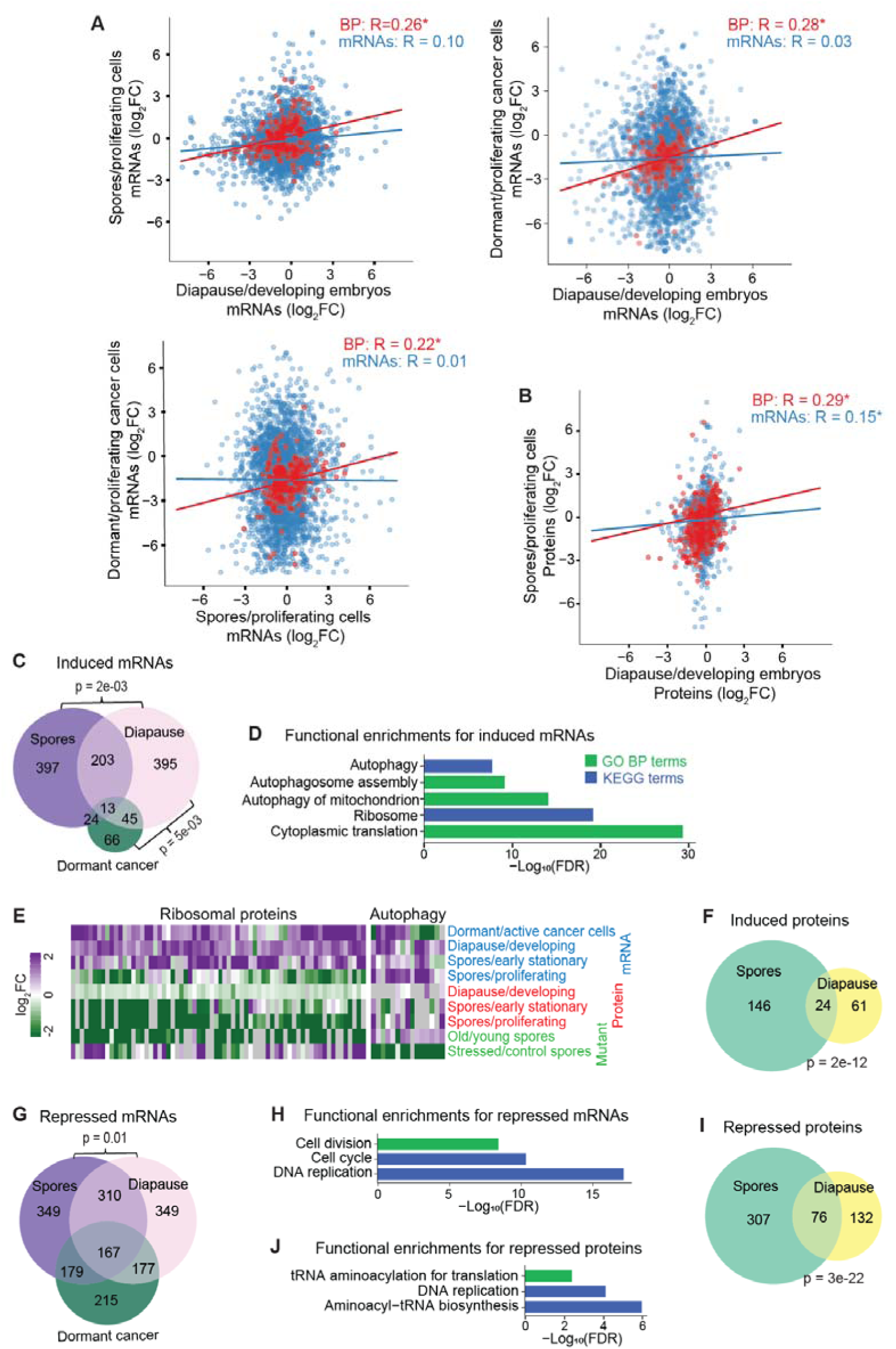
Genome regulation in yeast spores, killifish diapause embryos and human dormant cancer cells. (**A**) Scatter plots comparing log_2_ fold-changes (FC) in mRNA levels of orthologous genes in dormant cells relative to active control cells. Top left: yeast spores vs killifish diapause embryos; top right: human dormant cancer cells vs killifish diapause embryos; bottom left: human dormant cancer cells vs yeast spores. Each point represents either an individual mRNA (blue) or the mean log□FC of all genes within a GO Biological Process (red). The Pearson correlation coefficients for individual mRNAs (blue) or Biological Processeses (BP, red) are indicated at the top right of each plot, with asterisks indicating significant correlations (p <0.05). (**B**) Scatter plot comparing the log_2_ fold-changes (FC) in protein levels of orthologous genes in yeast spores relative to proliferating cells and in killifish diapause embryos relative to developing embryos. Each point represents either an individual protein (blue) or the mean log□FC of all genes within a GO Biological Process (red). The Pearson correlation coefficients for individual mRNAs (blue) or Biological Processeses (BP, red) are indicated at top right, with asterisks indicating significant correlations (p <0.05). (**C**) Overlaps among mRNAs induced (fold-change >1.5x; FDR <0.05) in yeast spores, killifish diapause embryos, and dormant cancer cells. In total, 2239 human genes with orthologs in killifish and yeast were detected in all 3 datasets. The p-values are indicated (Fisher’s exact test) for the overlaps that are higher than expected by chance. (**D**) Functional enrichments for mRNAs induced in at least 2 of the 3 dormant states. Green: GO Biological Processes; blue: KEGG pathways. The x-axis displays the –log_10_ of the false discovery rate (FDR), indicating the significance of the enrichment, determined using g:Profiler2. (**E**) Expression changes of conserved genes (columns) encoding cytoplasmic ribosomal proteins and autophagy proteins across the different datasets (rows) as labelled at right, including data for mRNAs (blue), proteins (red), and mutants (brown). Data for 58 ribosomal genes and 21 autophagy genes conserved in yeast, killifish, and human are shown, with log_2_ FC color-coded as indicated at left. (**F**) Overlap among proteins induced (fold-change >1.5x; FDR <0.05) in yeast spores and killifish diapause. In total, 1103 orthologous proteins were detected in both species. The p-value is indicated for the overlap (Fisher’s exact test). (**G**) Overlaps among mRNAs repressed (fold-change >1.5x; FDR <0.05) in yeast spores, killifish diapause, and dormant cancer cells. The p-value is indicated for the overlap (Fisher’s exact test). (**H**) Functional enrichments for mRNAs that are repressed in at least 2 of the 3 dormant states, as in (D). (**I**) Overlap among proteins repressed (fold-change >1.5x; FDR <0.05) in yeast spores and killifish diapause. The p-value is indicated for the overlap (Fisher’s exact test). (**J**) Functional enrichments for proteins that are repressed in both spores and killifish diapause, as in (D).

To define a core expression signature for dormant cells, we looked for genes regulated in the same direction in yeast spores, killifish diapause embryos, and human dormant cancer cells. For induced mRNAs, spores and diapause embryos showed the most overlap (216 genes), followed by diapause embryos and dormant cancer cells (58 genes), and spores and dormant cancer cells (37 genes) (Figure 3C; Dataset S3). A set of 13 genes was induced in all 3 types of dormant cells (Figure 3C), including 3 genes with autophagy-related functions, 3 genes involved in protein translation, and 2 genes with transcriptional roles (Dataset S3). We detected 55 priority unstudied genes across all 3 datasets, of which 15 were induced in at least 2 of the 3 dormant states (Dataset S3). The genes induced in dormant cells of at least 2 organisms were enriched for autophagy and protein translation (Figure 3D). Notably, mRNAs encoding autophagy and ribosomal proteins are also induced in dormant dauer larvae of worms (Boeck et al. 2016). Most conserved ribosomal-protein genes in yeast, killifish and humans were induced as mRNAs but repressed as proteins in all types of dormant cells analyzed, while most conserved autophagy genes were induced as both mRNAs and proteins (Figure 3E). Moreover, most mutants in these ribosomal and autophagy genes had antagonistic spore phenotypes with respect to longevity and stress resilience (Figure 3E). These results highlight that ribosomal and autophagy genes are core expression markers in dormant cells, where they play critical roles, as shown by our genetic results in yeast spores.

Among the detected proteins, 24 were induced in both spores and diapause embryos, significantly more than expected by chance (Figure 3F; Dataset S3). These proteins function in cellular detoxification and homeostasis, cytoskeleton organisation, chromatin organisation, and carbohydrate metabolism (Dataset S3). Four genes of different functions were induced at both the mRNA and protein levels in spores and diapause embryos, including SORD and GALE (carbohydrate metabolism), SORT1 (vesicle-mediated transport), and ATP2A3 (ion homeostasis).

For repressed mRNAs, spores and diapause embryos again showed the most overlap (477 genes), followed by spores and dormant cancer cells (346 genes) and diapause embryos and dormant cancer cells (344 genes), with 167 genes being repressed in all 3 types of dormant cells (Figure 3G; Dataset S3). These genes were enriched for cell proliferation and DNA replication (Figure 3H). Among the 55 priority unstudied genes, 26 were repressed in at least 2 of the 3 species, and 3 were repressed in all 3 species (Dataset S3). The repressed proteins, on the other hand, showed a more significant overlap, with 76 proteins repressed in both spores and diapause embryos (Figure 3I; Dataset S3). These proteins were enriched for DNA replication and protein translation, including aminoacyl-tRNA synthetases whose repression could contribute to ribosome pausing and decoupling mRNAs and proteins (Figure 3J). Thirty-three genes, enriched for DNA replication, were repressed at both the mRNA and protein levels in spores and diapause embryos (Dataset S3).

We compared our core genes to a published ‘G0 arrest transcriptional signature’ derived from cell-cycle arrested quiescent cells in primary solid tumors (Wiecek et al. 2023). The 2 gene lists shared many of the repressed genes, while only 1 induced gene, *SERINC1,* was shared (Dataset S3). This limited overlap could reflect differences between dormant and G0 cells. In G0 cells, a dormant-like state was induced by external factors that halt the cell cycle (e.g., serum starvation, contact inhibition), which might not fully reflect the physiological dormant states observed in cancer and other contexts studied here. The overlap in repressed genes could reflect a general inhibition of cell-cycle processes in non-dividing cells, such as DNA replication.

Together, these analyses reveal core biological processes whose gene-expression regulation is evolutionarily conserved across diverse dormant states and species, most notably the induction of autophagy– and translation-related processes and the repression of cell cycle-related processes. Commonly regulated genes may reflect conserved regulatory mechanisms. Indeed, a TFEB–TGFβ signaling axis controls adult reproductive diapause in worms, embryonic stem cell longevity in mice, and cancer diapause in humans (Nonninger et al. 2025). Process-level regulation is more conserved than gene-level regulation, highlighting the value of process-centric approaches. This finding is consistent with comparative studies of gene regulatory networks, revealing that evolutionary selection may conserve the control of pathways or processes rather than that of particular genes within them (D. Thompson et al. 2015). Nevertheless, we identified a few core genes whose mRNAs are universally induced or repressed across yeast spores, killifish diapause embryos and human dormant cancer cells. These overlaps are most significant between yeast spores and killifish diapause embryos and are even more pronounced at the protein level.

### Transcriptomes and proteomes change in old and stressed yeast spores

We next investigated whether dormant yeast spores can modulate genome expression with prolonged time or in response to an external cue. To this end, we analysed the transcriptomes and proteomes of spores stored for 2 weeks, 3 months, and 5 months, as well as 60 minutes after a 45°C heat shock (Figure 4A, top half). We detected 3986 mRNAs (78.8%), 2650 proteins (52.4%), and 1896 long non-coding RNAs (24.9%) in at least 5 samples. The transcriptomic signatures showed substantial variance between these datasets (Figure 4B): the samples from aged spores (3 and 5 months) were similar to each other but distinct from the young samples (2 weeks), whereas samples exposed to heat shock were more similar to the young samples than aged spores.

**Figure 4:**
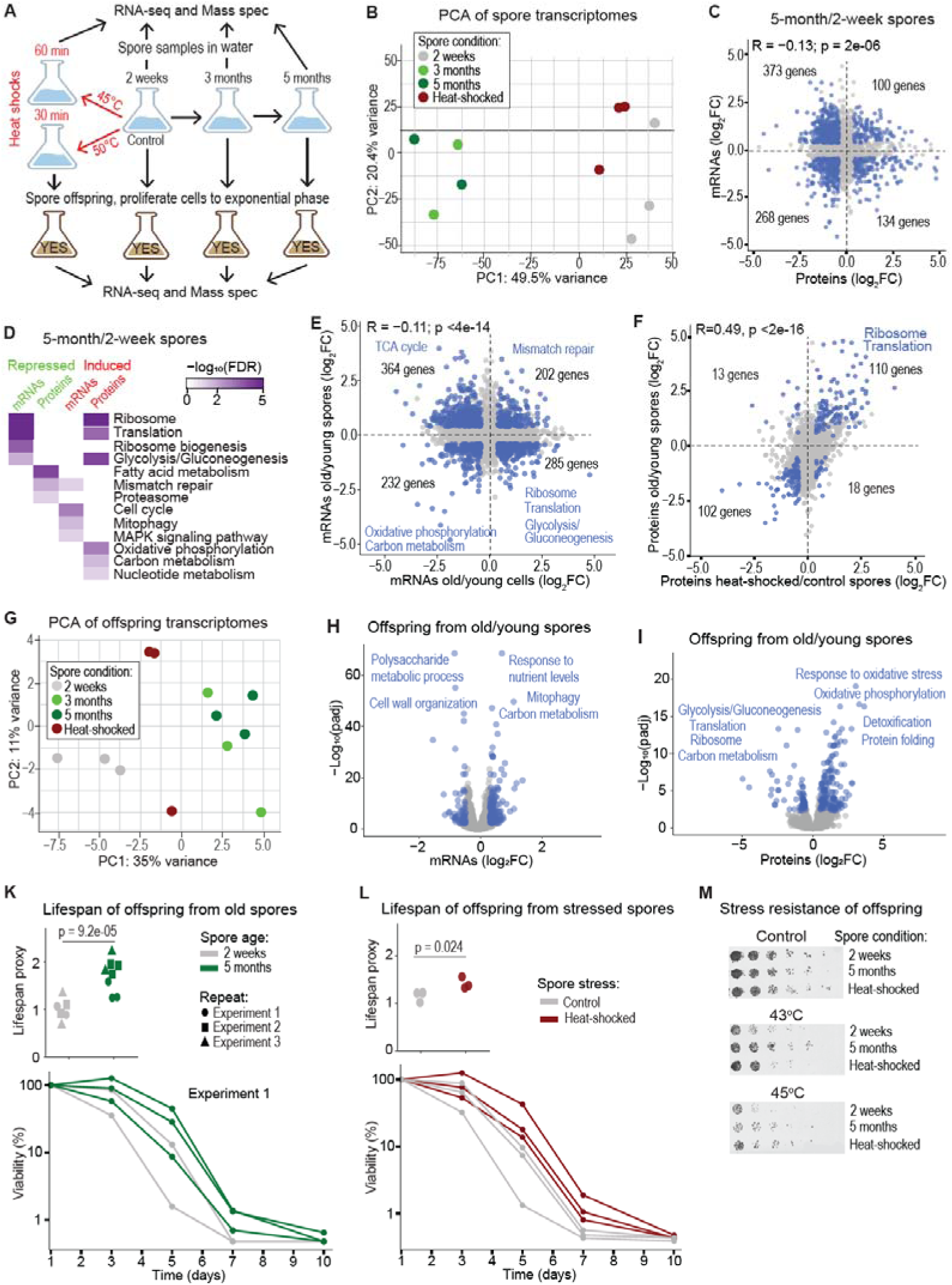
Transcriptome and proteome changes in older and stressed spores and their offspring. (**A**) Scheme of the experimental design. Spores were aged and heat-shocked as indicated, followed by transcriptome and proteome analyses of the different spore samples (top) and of the offspring cells derived from germinating the different spore samples and growing for ∼18 hours to exponential phase (bottom). For the 2-week-old spores, with and without heat shock, 3 biological replicates were analysed for each condition, whereas for the 3– and 5-month-old spores, 2 biological replicates were analysed. For all offspring cells, 3 biological replicates were analysed for each condition. **(B)** Principal Component Analysis (PCA) of transcriptomic signatures from spore samples after different times or heat shock. Each dot represents a biological replicate, and the colors correspond to the conditions as indicated. The variances represented in the principal components 1 and 2 are indicated on the axes. The gene expression values used for the PCA are normalized read counts derived from DESeq2. **(C)** Scatterplot comparing the log_2_ fold-changes (FC) in mRNA and protein levels in 5-month-old relative to 2-week-old spores. The Pearson correlation coefficient (R) and its significance are displayed on top. Genes differentially expressed (FC >1.5x; FDR <0.05) at both the mRNA and protein level are highlighted in blue, and their numbers are indicated for each quadrant. **(D)** Enriched GO Biological Processes and KEGG pathways associated with repressed (green) and induced (red) mRNAs and proteins as indicated in 5-month-old compared to 2-week-old spores. The colour shade corresponds to the false discovery rate (FDR; top right). Enrichments were calculated with g:Profiler2. **(E)** Scatterplot showing the log_2_ FC in mRNA levels in 5-month-old vs 2-week-old spores compared to old vs young stationary-phase cells at 100% and 50% viability (Atkinson et al. 2018). The Pearson correlation coefficient and its significance are shown on top. Genes differentially expressed (FC >1.5x; FDR <0.05) in both spores and cells are highlighted in blue, with other genes in grey. The number of significant genes along with enriched GO and KEGG terms is indicated for each quadrant (g:Profiler2). **(F)** Scatterplot showing the log_2_ FC in protein levels in 5-month-old vs 2-week-old spores compared to heat-shocked vs control spores. The Pearson correlation coefficient and its significance are shown on top. Proteins differentially expressed (FC >1.5x; FDR <0.05 in at least one condition) in both old spores and heat-shocked spores are highlighted in blue, with other proteins in grey. The number of significant proteins, along with enriched GO terms (g:Profiler2), is indicated for each quadrant. **(G)** PCA of transcriptomic signatures from offspring cells derived from spores after different times or heat shock. Each dot represents a biological replicate, and the colors correspond to the spore conditions as indicated. The variances represented in the principal components 1 and 2 are indicated on the axes. The gene expression values used for the PCA are normalized read counts derived from DESeq2. **(H)** Volcano plot showing mRNA expression in offspring cells from 5-month-old spores relative to offspring from 2-week-old spores. Blue dots: genes that are significantly lower (72 genes) or higher (160 genes) expressed in the offspring from old spores, with other genes in grey. Enriched GO and KEGG terms for induced and repressed genes are indicated. Significance thresholds for differential expression: FC >1.32x; FDR <0.05. **(I)** Volcano plot showing protein expression in offspring cells from 5-month-old spores relative to offspring from 2-week-old spores. Blue dots: proteins that are significantly lower (65 proteins) or higher (158 proteins) expressed in the offspring from old spores, with other proteins in grey. Enriched GO and KEGG terms for induced and repressed proteins are indicated. Significance thresholds for differential expression: FC >1.32x; FDR <0.05. (K) Top: Normalized chronological lifespan proxies for cells derived from young (grey) and old (green) spores, with the p-value for the difference in lifespan indicated (T-test). Cells germinated from spores in YES medium were grown to stationary phase, diluted and grown again to stationary phase, with Day 1 being the time when cells reached maximum density. The lifespan proxies represent the time required for cells derived from old spores to reach 1% viability, relative to those from young spores. The lifespans were measured for 6 (young spores) and 8 (old spores) independent repeats across 3 experiments, using a robotics-based colony-forming unit assay (Romila et al. 2021), with values normalized to Day 1 (100% viability). Bottom: Chronological lifespan curves for Experiment 1, including 2 independent repeats for offspring from young spores (grey) and 3 repeats for offspring from old spores (green). The lifespan curves of Experiments 2 and 3 are provided in Figure S3E. (**L**) Top: Normalized chronological lifespan proxies for cells derived from control spores (grey) and spores heat-shocked for 30 minutes at 50°C (red), determined as in (K), with the p-value for the difference in lifespan indicated (T-test). Bottom: Chronological lifespan curves for this experiment, including 3 independent repeats for offspring from control spores (grey) and 3 repeats for offspring from heat-stressed spores (red). (**M**) Spot test of cells derived from young, old, and heat-shocked spores. Cells germinated from spores in YES medium were grown to stationary phase, diluted and grown again to stationary phase. Cells were 3-fold serially diluted and spotted onto YES plates. To test for stress resistance, cells were incubated at 43°C or 45°C for 30 min before plating.

**Figure S3:**
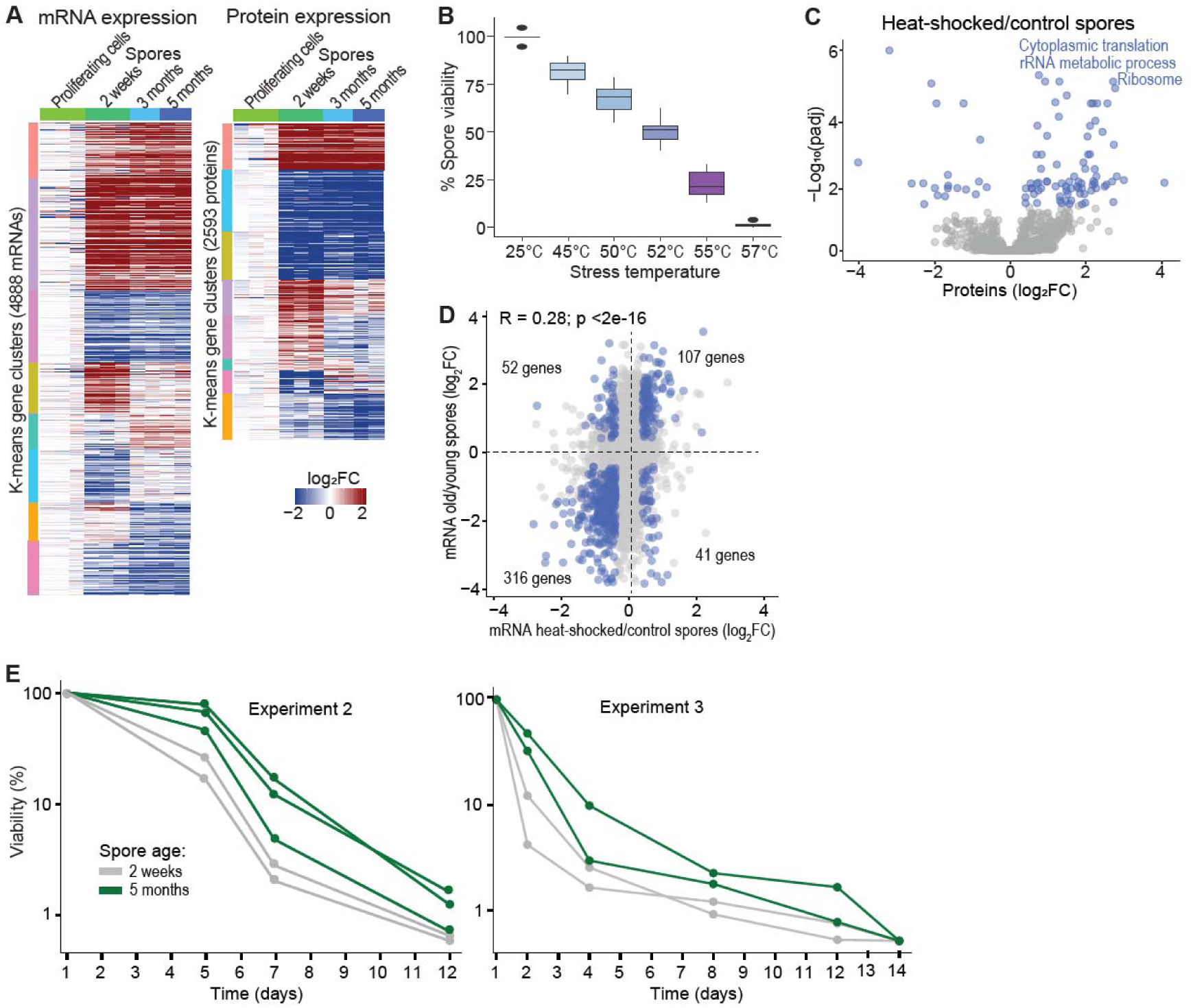
Transcriptome and proteome changes in older and stressed spores and their offspring. (**A**) Heatmaps of transcriptome (left) and proteome (right) changes in spores after 2 weeks, 3 months, and 5 months. Rows represent genes, and the colors show the fold-changes (FC) in expression relative to proliferating cells. Columns represent the different conditions, with data for the 2 to 3 biological repeats provided separately. Only genes that are differentially expressed (FC >1.5x; FDR <0.05) in at least 1 condition are shown. Gene expression patterns are grouped into 6 clusters using k-means. (**B**) Boxplot showing the percentage viability of 2-week-old spores following a 30-minute heat-shock treatment at the indicated stress temperatures. (**C**) Volcano plot showing protein expression in heat-shocked relative to control spores. Blue dots: proteins that are significantly lower or higher expressed in heat-shocked spores. Enriched GO terms for induced proteins are indicated. Significance thresholds for differential expression: FC >1.5x; FDR <0.05. (**D**) Scatterplot comparing the log_2_ fold-changes (FC) in mRNA levels in 5-month-old vs 2-week-old spores compared to mRNA levels in heat-shocked vs control spores. The Pearson correlation coefficient (R) and its significance are displayed on top. Genes differentially expressed (FC >1.5x; FDR <0.05) in both old and heat-shocked spores are highlighted in blue, and the numbers of these genes are indicated for each quadrant. (E) Chronological lifespan curves of Experiments 2 and 3 for offspring from young (grey) and old (green) spores for the results shown in Figure 3K.

To examine gene-expression changes in older spores, we performed k-means clustering on genes differentially expressed at any time point relative to proliferating cells. About half of the regulated transcripts and proteins showed altered expression in older spores (3 and 5 months) compared to 2 weeks (Figure S3A). Accordingly, 1838 mRNAs, 740 proteins, and 593 long non-coding RNAs were differentially expressed in 5-month-old spores relative to 2-week-old spores (Dataset S4). Notably, the mRNA and corresponding protein expression changes were slightly inversely correlated (Figure 4C), suggesting substantial regulation at post-transcriptional levels leading to decoupled transcript and protein expression. Such decoupled gene expression also features in quiescent stem cells and during aging and associated diseases (Dick et al. 2023; Takemon et al. 2021; Janssens et al. 2015; de Morree and Rando 2023). Accordingly, functional enrichments differed between mRNAs and proteins whose levels changed in old relative to young spores (Figure 4D). Most notably, ribosome– and translation-related processes became strongly repressed at the mRNA level but induced at the protein level (Figure 4D). These changes might reflect the importance of translational capacity for spore germination once environmental conditions allow it. In starving budding yeast cells, a pool of ribosomes is maintained in an inactive state, required to restart cell growth when conditions allow (Van Dyke et al. 2013; Shetty et al. 2023). Similarly, in starved fission yeast cells, ribosomes ‘hibernate’ on mitochondria whilst protein synthesis is suspended (Gemin et al. 2024). Older spores also showed a relative increase in carbon metabolism and mitochondrial processes at the protein level (Figure 4D). We conclude that the relative levels of many RNAs and proteins in spores change with time and thus differ between young and old spores. These changes are not coordinated between RNAs and corresponding proteins, leading to decoupled expression.

We wondered whether the expression changes in older spores are similar to those in chronologically aging cells of *S. pombe* and thus reflect an aging signature. To investigate this, we compared transcriptomic changes in old vs young spores with those in old vs young stationary-phase cells (Atkinson et al. 2018). Notably, the changes in old spores and old cells showed a weak inverse correlation (Figure 4E). Key processes such as translation, the TCA cycle, and glycolysis/gluconeogenesis were regulated in opposite directions in old spores and old cells, whereas other carbon– and energy-metabolism processes and mismatch repair were regulated in the same direction (Figure 4E). These results indicate that the transcriptome changes observed over time in spores are distinct from those in aging cells, which also show much shorter lifespans than spores.

To test whether dormant spores can still respond to environmental stress, we subjected them to a heat shock at 45°C after two weeks (Figure 4A). This temperature killed <25% of spores (Figure S3B). Compared to old spores, much fewer transcripts and proteins were differentially expressed in heat-shocked spores relative to control spores: 151 mRNAs, 63 proteins, and 71 long non-coding RNAs (Dataset S4). The mRNA and corresponding protein expression changes in stressed spores were not correlated (R = 0.02; p = 0.46). As in old spores, proteins induced in stressed spores were enriched for translation-related processes (Figure S3C). The genes differentially expressed in both old and stressed spores were correlated, both at the transcriptomic level (Figure S3D) and, even more so, at the proteomic level (Figure 4F). We conclude that dormant spores change their transcriptome and proteome with time and in response to environmental stress. The expression changes in old spores are similar to those in stressed spores, albeit the latter involve many fewer genes, but they differ from those in aging cells. These patterns suggest that changes in spore expression, including the induction of translation-related proteins, may reflect active regulatory processes that support long-term viability.

Our findings indicate that fission yeast spores can tune their transcriptomes and proteomes in response to the passage of time or to changing environments. These results are consistent with previous work that has suggested that fungal dormancy is not a static state: in budding yeast spores, the maintenance of gene expression is critical for their survival (Maire et al. 2020), and fission yeast spores are affected by both environmental conditions and time of dormancy, altering spore fitness like stress tolerance and viability (Nuckolls et al. 2025). Similarly, the transcriptome of bacterial spores is dynamic and affected by age or environmental changes (Segev et al. 2012).

### Spore age and stress experience affect gene expression and cellular traits in cells derived from these spores

We then investigated whether offspring cells from aged or stressed yeast spores showed any lasting changes in their transcriptome or proteome. We analyzed gene expression in cells derived from spores stored for 2 weeks, 3 months, and 5 months, or heat-shocked at 50°C for 30 minutes; all offspring were obtained by germinating spores in rich medium and growing the cells to mid-exponential phase for ∼6 generations (Figure 4A, bottom half). A PCA revealed substantial differences in the transcriptome signatures among these datasets (Figure 4G): cells derived from aged spores (3 and 5 months) were similar to each other but distinct from cells derived from young spores (2 weeks), while the heat shock had a smaller effect than prolonged time. These differences were qualitatively similar to those in the corresponding spore samples (Figure 4B). Accordingly, several transcripts and proteins were differentially expressed in offspring from 5-month-old spores relative to offspring from 2-week-old spores, including 199 mRNAs, 223 proteins, and 33 long non-coding RNAs (Dataset S5). The mRNA and corresponding protein expression changes in offspring from old vs young spores were not correlated (R = 0.02; p = 0.50). Although many more genes were differentially expressed in aged spores than in the corresponding offspring cells, the offspring showed clear expression signatures with functional enrichments at both the transcriptomic (Figure 4H) and proteomic (Figure 4I) levels. Differentially expressed genes were enriched for some similar biological processes in spores and offspring, including those related to carbon metabolism, respiration, protein translation, mitophagy, and stress responses, although the processes were sometimes regulated in opposite directions in spores and offspring, especially at the protein level (Figure 4D,H,I). Overall, the gene-expression changes in old spores and their corresponding offspring cells were only weakly correlated at the transcriptomic and proteomic levels (R = 0.16; p <2e-16 in both cases). On the other hand, gene-expression changes in offspring cells from old spores showed moderate correlations with those from heat-shocked spores at the transcriptome and proteome levels (R = 0.56 and 0.23, respectively; p <2e-16). Although we cannot exclude rare genetic changes or clonal selection as contributors to the persistent expression differences in offspring cells, this seems unlikely given the similar outcomes of independent biological repeats (Figure 4G). We conclude that spore age and heat-stress experience have similar long-term effects on gene expression in cells derived from these spores, even after several generations of proliferation.

We wondered whether spore age or heat stress also affects cellular phenotypes of offspring derived from these spores. To address this question, we determined the chronological lifespan and stress resistance for offspring cells from different spore samples. Remarkably, cells derived from 5-month-old spores showed longer chronological lifespans during stationary phase than cells derived from 2-week-old spores (Figure 4K; Figure S3E). Similarly, cells derived from heat-shocked spores showed longer lifespans than those derived from unstressed control spores (Figure 4L). These lifespan effects were modest but reproducible and significant. Moreover, offspring cells derived from older or heat-shocked spores exhibited higher resistance to heat stress (Figure□4M). We conclude that spore age and heat-stress experience also have long-term effects on cellular phenotypes in the offspring from these spores, including a prolonged chronological lifespan and increased heat resistance. Thus, cells derived from old or stressed spores ‘remember’ these states across spore germination and for several generations of cell divisions.

### Transcriptomes and proteomes change in older and stressed killifish diapause embryos

We investigated whether killifish diapause embryos, like yeast spores, modulate their transcriptome or proteome with prolonged time or in response to a heat shock. We analyzed gene expression in diapause embryos after 2 weeks (early diapause) and 3 months (late diapause), as well as 60 minutes after a heat shock in early diapause embryos (Figure 5A). In total, among the 26,141 annotated genes, we detected 16,178 mRNAs (61.9%) and 1,729 proteins (6.6%) in at least 3 samples. As in spores, the different conditions showed distinct transcriptomic profiles (Figure 5B). Unlike in spores (Figure 4B), heat shock had a larger effect on mRNA expression than prolonged time in diapause embryos (Figure 5B).

**Figure 5:**
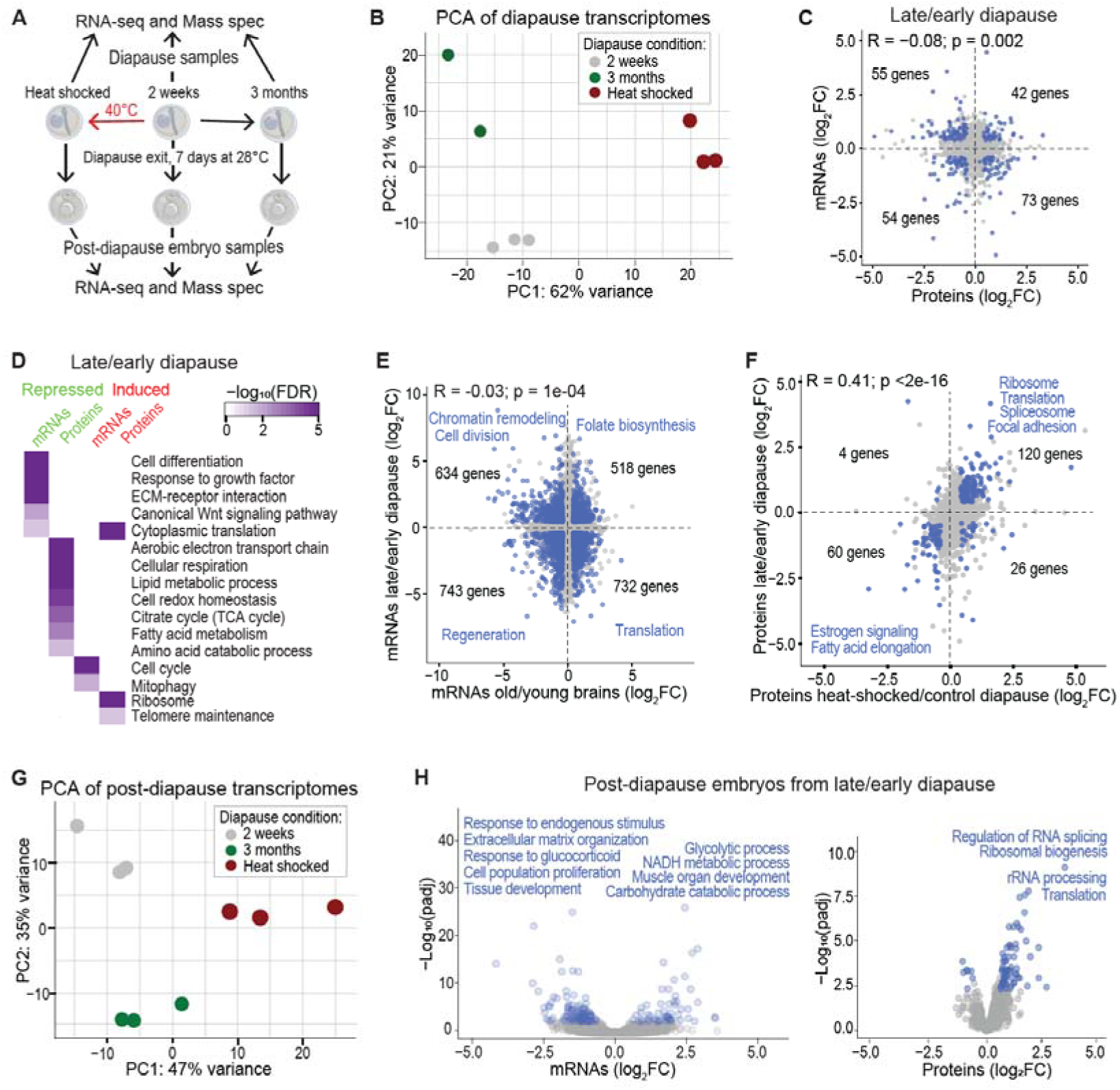
Transcriptome and proteome changes in late and stressed diapause and post-diapause embryos. (**A**) Scheme of the experimental design. *N. furzeri* diapause embryos (MZCS strain) were maintained for 2 weeks (early, with or without heat shock) and 3 months (late) as indicated, followed by transcriptome and proteome analyses of the different diapause samples (top) and the post-diapause embryos derived from the diapause samples by triggering diapause exit at increased temperature and development for one week to the prehatching stage (bottom). For the early diapause samples, 3 biological replicates were analysed; for late diapause, 2 and 3 biological replicates were analysed for transcriptomics and proteomics, respectively; for all post-diapause samples, 3 biological replicates were analysed. For diapause samples, batches of ∼50 and ∼15 embryos were used for transcriptomics and proteomics, respectively; for post-diapause samples, batches of ∼10 and ∼ 1 embryos were used for transcriptomics and proteomics, respectively. **(B)** Principal Component Analysis (PCA) of transcriptomic signatures from early, late, and heat-shocked diapause samples. Each dot represents a biological replicate, and the colors correspond to the conditions as indicated. The variances represented in the principal components 1 and 2 are indicated on the axes. The gene expression values used for the PCA are normalized read counts derived from DESeq2. **(C)** Scatterplot comparing the log_2_ fold-changes (FC) in mRNA and protein levels in 3-month-old relative to 2-week-old diapause embryos. The Pearson correlation coefficient (R) and its significance are displayed on top. Genes differentially expressed (FC >1.5x; FDR <0.05) at both the mRNA and protein level are highlighted in blue, and their numbers are indicated for each quadrant. **(D)** Enriched GO Biological Processes and KEGG pathways associated with repressed (green) and induced (red) mRNAs and proteins as indicated in 3-month compared to 2-week diapause embryos. The colour shade corresponds to the false discovery rate (FDR; top right). Enrichments were calculated with g:Profiler2. **(E)** Scatterplot showing the log_2_ FC in mRNA levels in 3-month vs 2-week diapause embryos compared to old vs young brain cells (Baumgart et al. 2014). The Pearson correlation coefficient and its significance are shown on top. Genes differentially expressed (FC >1.5x; FDR <0.05) in both diapause and brain cells are highlighted in blue, with other genes in grey. The number of significant genes and enriched GO/KEGG terms are indicated for each quadrant (g:Profiler2). **(F)** Scatterplot showing the log_2_ FC in protein levels in 3-month vs 2-week diapause embryos compared to heat-shocked vs control embryos. The Pearson correlation coefficient and its significance are shown on top. Proteins differentially expressed (FC >1.32; FDR <0.05 in ≥1 condition) in both late and heat-shocked diapause embryos are highlighted in blue, with other proteins in grey. The number of significant proteins, along with enriched GO terms (g:Profiler2), is indicated for each quadrant. **(G)** PCA of transcriptomic signatures from post-diapause embryos derived from early, late, or heat-shocked diapause. Each dot represents a biological replicate, and the colors correspond to the spore conditions as indicated. The variances represented in the principal components 1 and 2 are indicated on the axes. The gene expression values used for the PCA are normalized read counts derived from DESeq2. **(H)** Volcano plot showing mRNA (left graph) and protein (right graph) expression in post-diapause embryos derived from 3-month vs 2-week diapause. Blue dots: genes that are significantly lower or higher expressed in the post-diapause embryos from 3-month diapause, with other genes in grey. Enriched GO and KEGG terms for induced and repressed genes are indicated. Significance thresholds for differential expression: FC >1.32x; FDR <0.05.

**Figure S4:**
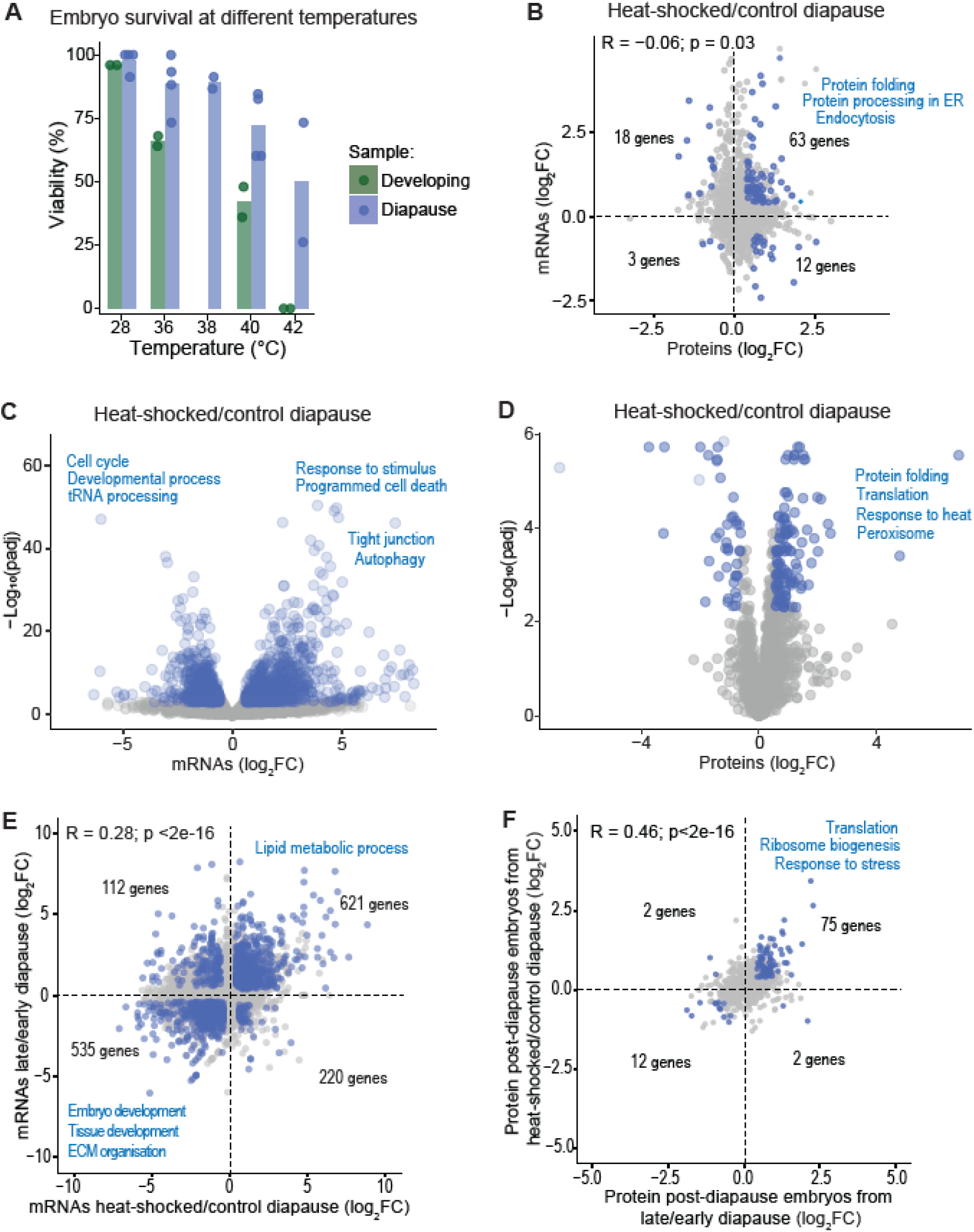
Transcriptome and proteome changes in late and stressed diapause and post-diapause embryos. (**A**) Viability of diapause embryos (blue) and developing control embryos (green) after a 1-hour heat-shock at the indicated temperatures. Survival was assessed visually 24 hours after the heat shock. Bars indicate the mean survival for each condition, and dots represent independent repeats. For developing samples, batches of 25 embryos were analysed, whereas for diapause samples, batches ranged from 15 to 23 embryos. **(B)** Scatterplot comparing the log_2_ fold-changes (FC) in mRNA and protein levels in heat-shocked relative to control diapause embryos. The Pearson correlation coefficient (R) and its significance are displayed on top. Genes differentially expressed (FC >1.5x; FDR <0.05) at both the mRNA and protein level are highlighted in blue, with other genes in grey. The number of significant genes and the enriched GO terms are indicated for each quadrant. **(C)** Volcano plot showing mRNA expression in heat-shocked relative to control diapause embryos. Blue dots: genes that are significantly lower or higher expressed after the heat shock. Enriched GO terms for repressed and induced mRNAs are indicated. Significance thresholds for differential expression: FC >1.5x; FDR <0.05. **(D)** Volcano plot showing protein expression in heat-shocked relative to control diapause embryos. Blue dots: proteins that are significantly lower or higher expressed after the heat shock. Enriched GO terms for induced proteins are indicated. Significance thresholds for differential expression: FC >1.5x; FDR <0.05. **(E)** Scatterplot showing the log_2_FC in mRNA levels in 3-month vs 2-week diapause embryos compared to heat-shocked vs control diapause embryos. The Pearson correlation coefficient and its significance are shown on top. Genes differentially expressed (FC >1.32x; FDR <0.05) in both conditions are highlighted in blue, with other genes in grey. The number of significant genes and the enriched GO terms are indicated in the quadrants. **(F)** Scatter plots comparing log_2_FC in protein levels in post-diapause embryos from heat-shocked vs control diapause compared to post-diapause embryos from 3-month vs 2-week diapause. The Pearson correlation coefficient (R) and its significance are displayed. Genes differentially expressed (FC >1.32x; FDR <0.05) in both conditions are highlighted in blue, with other genes in grey. The number of significant genes and the enriched GO terms are indicated for each quadrant.

In late relative to early diapause embryos, 708 mRNAs and 395 proteins were differentially expressed (Dataset S6). As in spores, the mRNA and corresponding protein expression changes were slightly inversely correlated (Figure 5C), suggesting substantial post-transcriptional regulation in both species. Consistently, functional enrichment analysis revealed differences between mRNAs and proteins whose levels changed in late diapause embryos (Figure 5D). For example, respiration-related genes were repressed only at the protein level and developmental functions, such as cell differentiation, growth-factor response and Wnt signaling, were only induced at the mRNA level. The latter enrichments suggest that late diapause embryos are inducing transcripts required to resume active development. Intriguingly, these enrichments showed some similarities to those in yeast spores, including cell-cycle and mitophagy genes being induced at the mRNA level, fatty-acid metabolism genes being repressed at the protein level, and translation-related genes being repressed at the mRNA level but induced at the protein level (Figure 4D, Figure 5D). Overall, slightly more mRNAs and proteins were regulated in both old spores and late diapause embryos than expected by chance (p = 0.037 and 0.048, respectively; Fisher’s exact test): 142 mRNAs (87 regulated in same direction) and 88 proteins (49 regulated in same direction). As for spores, we conclude that the relative levels of many transcripts and proteins differ between early and late diapause embryos, but these changes over time are not coordinated between RNAs and their corresponding proteins, as reflected by a loss of expression correlation.

We then checked whether the expression changes in late diapause embryos are similar to the transcriptome and proteome changes reported during brain aging in adult killifish (Kelmer Sacramento et al. 2020; Baumgart et al. 2014). We found weak inverse correlations in expression changes for both the transcriptomes (Figure 5E) and proteomes (R_Pearson_ = – 0.004). This result mirrors the inverse correlation between expression changes in older spores and aging cells of fission yeast (Figure 4E). Processes like chromatin remodelling, cell division, and translation were regulated in opposite directions in old brains compared to late diapause embryos, the latter mirroring the results in fission yeast (Figure 4E, Figure 5E). We conclude that the gene-expression changes observed in late diapause embryos are distinct from an aging signature in brain cells. This finding suggests that, unlike adult killifish, embryos may not age during diapause, consistent with results showing that time spent in diapause does not negatively affect subsequent adult growth, fertility, or lifespan (Hu et al. 2020). Instead, the observed expression changes in late diapause could reflect measures to maintain long-term viability and prepare for re-initiating active development.

To test whether diapause embryos can respond to environmental stress, we subjected early diapause embryos to 1 hour at 40°C. This temperature was chosen as a robust stress that maintains ∼60-80% survival in diapause embryos, whereas actively developing embryos showed less than 50% survival (Figure S4A). As elevated temperature can trigger diapause exit, we returned the heat-shocked embryos to 20°C and monitored them for 3 days, revealing that all embryos remained in diapause after the transient heat shock. In heat-shocked relative to control diapause embryos, 1739 mRNAs and 157 proteins were differentially expressed (Dataset S6). The mRNA and corresponding protein expression changes in stressed embryos were marginally inversely correlated (Figure S4B). This result indicates that, as in yeast spores, the stress response at the transcriptomic level is decoupled from that at the proteomic level. Despite this overall lack of correlation, genes involved in protein folding, endocytosis, and endoplasmic reticulum processing were induced at both the mRNA and protein levels in response to heat (Figure S4B). Induced mRNAs were also enriched for processes like autophagy, stimulus response, and programmed cell death, while repressed mRNAs were enriched for cell-cycle and developmental processes (Figure S4C). These findings are consistent with results from diapause embryos of the killifish *Austrofundulus limnaeus,* which regulate similar processes at the transcriptome level in response to hypoxia stress (Clouser et al. 2025). On the other hand, the proteins induced after the heat shock were enriched for processes related to translation, peroxisomes, and the heat response (Figure S4D). As in spores, the genes differentially expressed in both late and stressed diapause showed correlated changes at the transcriptome level (Figure S4E) and, even more so, at the proteome level (Figure 5F). This finding mirrors the patterns observed for yeast spores (Figure S3D, Figure 4F). Moreover, in both spores and diapause embryos, ribosome– and translation-related processes were enriched among the proteins induced by both stress and prolonged time (Figure 4E, Figure 5F).

### Diapause duration and stress experience affect gene expression in post-diapause embryos

As in yeast, we then investigated whether developing post-diapause killifish embryos exposed to different diapause conditions showed any lasting changes in their transcriptomes or proteomes. We analyzed gene expression in post-diapause embryos at the pre-hatching stage, obtained from early, late or stressed diapause embryos (Figure 5A, bottom half). The different post-diapause (prehatching) embryos showed distinct transcriptome signatures (Figure 5G). Accordingly, 160 to 210 transcripts and 77 to 127 proteins were differentially expressed in prehatching embryos derived from late or stressed diapause embryos relative to those derived from early diapause embryos (Dataset S7). The mRNA and corresponding protein expression changes in prehatching embryos from late vs early diapause were marginally inversely correlated (R = −0.082; p = 0.002). As in yeast, many more genes were differentially expressed in late vs early diapause than in the corresponding prehatching embryos; the latter showed clear expression signatures with functional enrichments at both the transcriptomic and proteomic levels (Figure 5H). The induced mRNAs were enriched for carbon metabolism, and the induced proteins were enriched for protein translation and splicing (Figure 5H). This expression signature showed some similarities to the canonical heat-shock response in zebrafish embryos, including the regulation of carbohydrate metabolism (Feugere et al. 2023). Overall, the gene-expression changes in late diapause embryos and the corresponding prehatching embryos were not correlated at either the transcriptomic (R = 0.02; p = 0.07) or proteomic (R = 0.05; p = 0.17) level. Similar to the corresponding results in yeast, the gene-expression changes in prehatching embryos from late diapause spores showed moderate correlation with those from heat-shocked diapause at both the transcriptomic and proteomic levels (R = 0.33 and 0.46, respectively; p <2e-16). At the protein level, processes related to protein translation and stress response were induced in prehatching embryos derived from both late and heat-stressed diapause (Figure S4F). Overall, 6 orthologous mRNAs and 3 proteins were regulated both in offspring cells from old spores and in post-diapause embryos from late diapause. We conclude that the diapause duration and stress experience exert long-term effects on gene expression in post-diapause embryos that have developed for 7 days and are ready to hatch. These memory effects across several cell divisions after dormancy exit mirror the results in offspring cells from yeast spores.

### Conclusions

Our integrated multi-omics profiling of gene function and expression in distinct types of dormant cells and cells derived from these dormant cells reveals core biological processes, common patterns, and surprising features of dormancy. We provide a genome-wide survey of genes affecting the longevity and heat-shock resilience of fission yeast spores. Long-lived deletion mutants, including those involving the TCA cycle and ribosomal proteins, tend to be stress-sensitive, while short-lived mutants, including those involving autophagy proteins, tend to be stress-resistant. This surprising finding points to a trade-off between processes supporting spore longevity and those supporting stress resilience. Gene deletions affecting lifespan in spores show little overlap with those affecting lifespan in chronologically aging stationary-phase cells, indicating that several biological processes underlying longevity in dormant cells differ from those in aging cells. Mitochondrial respiration is important for the longevity of both spores and stationary-phase cells, while ribosome– and autophagy-related functions show opposite effects on the lifespans of spores and cells.

Both yeast spores and killifish diapause embryos feature substantially remodelled transcriptomes and proteomes compared to non-dormant reference cells. We observed a consistent trend of decoupled regulation of ribosomal proteins, with induction at the mRNA level and repression at the protein level across the two dormant states. This finding suggests an anticipatory post-transcriptional regulatory strategy where cells stabilize translation-related transcripts to facilitate a rapid return to growth upon dormancy exit or a rapid response to environmental challenges (see below). Many unstudied genes conserved from fission yeast to humans were differentially expressed in yeast spores and in killifish diapause, consistent with dormancy being an understudied cellular state.

Although the overlap between genes differentially expressed in spores and genes functioning in spore lifespan or heat resilience is not significant, processes related to protein translation, the mitochondrial TCA cycle, and autophagy are enriched among both gene sets. For example, autophagy-related genes are induced in spores and crucial for their heat resilience but detrimental for their longevity. These results illustrate that gene expression profiles are insufficient to infer functional relevance, and the multi-layered approach integrating transcriptomics, proteomics, and functional genomics applied here provides more nuanced insights.

A comparison of gene expression in yeast spores, killifish diapause embryos and human dormant cancer cells reveals that the regulation of specific genes often diverges, whereas the regulation of biological processes shows substantial conservation across the diverse dormant states and organisms. At the mRNA level, the induction of autophagy– and translation-related processes and the repression of cell cycle-related processes define core expression features of the dormant states examined here.

The relative levels of many transcripts and proteins change over time or after a heat shock in both yeast spores and killifish diapause embryos, indicating that dormancy is not a static state of ‘suspended animation’ but a dynamic, responsive state. Both species showed similar patterns of expression changes during dormancy, including decoupled mRNA and protein changes, similar responses to prolonged time and heat stress, and shared functional enrichments among the regulated genes. The uncoupled changes in mRNA and protein levels point to post-transcriptional processes. Notably, the transcriptome changes observed over time in spores or diapause show modest inverse correlations with those in chronologically aging cells in yeast or aging brain cells in killifish, respectively. This finding suggests that the expression changes in older dormant cells do not reflect aging-associated decline but active regulatory processes maintaining long-term viability, consistent with dormant cells featuring exceptionally long lifespans and being more resilient to aging. For example, ribosomal and other translation-related proteins are induced in stressed or older dormant cells, suggesting that the corresponding transcripts, which are present at high levels, are translated on demand to support a rapid stress response or prepare for dormancy exit, respectively. Remarkably, spore age and heat-stress exposure exert persistent effects on gene expression and cellular traits in the cells derived from these spores, including a prolonged chronological lifespan and increased heat resistance. Similarly, diapause duration and stress exposure exert long-term effects on gene expression in the corresponding post-diapause embryos at the pre-hatching stage. Thus, reminiscent of hormetic priming, both yeast and killifish dormant cells transmit molecular memories of their dormancy duration or stress experience to their offspring cells that have exited dormancy and undergone several generations of cell divisions. We propose that this memory functions through chromatin modifications, and further studies will be required to dissect the mechanistic basis of this epigenetic phenomenon.

We acknowledge some limitations of the study. Bar-seq quantification of deletion mutants was performed after spore germination and regrowth; thus, the barcode abundance reflects the combined effects of spore survival, germination efficiency, and growth, which may influence stress– or lifespan-associated phenotypes for some mutants, a compromise necessary to prevent barcodes of dead spores from being sequenced. As is the case for most expression studies, our transcriptome and proteome data reflect relative expression levels between two conditions. It is likely that the absolute cellular numbers of most transcripts and proteins are reduced in dormant cells, as they are in quiescent *S. pombe* cells (Marguerat et al. 2012). Thus, genes induced in dormant cells relative to active cells might be repressed in absolute levels, but to a lesser extent than most other genes. The proteome depth differs between yeast and killifish, and is lower than the transcriptome depth, limiting our comparisons at the protein level, particularly for low-abundance proteins. Finally, some of the expression changes detected in dormant cells may reflect passive processes, such as transcript or protein half-lives, rather than active regulatory responses. Nevertheless, even such passive changes could have biological effects.

## Materials and Methods

### Yeast strains and experimental conditions

Leupold’s laboratory strain *968 h^90^* (JB50) was used. For exponentially growing vegetative control cells, we used YES medium (yeast extract with supplements), and for stationary-phase control cells, we used YES agar plates. Mating, meiosis, and sporulation were induced on Malt Extract Agar (MEA) plates. For the transcriptomic and proteomic experiments, JB50 cells were grown in YES at 32°C overnight as a preculture. The cells were then spread onto MEA plates and incubated at 25°C for 5 days. After incubation, spores were harvested and washed 3 times with distilled water to remove any residual nutrients. Sporulation efficiency was assessed by light microscopy (Leica, MDG33) to observe spore formation. Spores were first treated with 30% ethanol for 30 min to eliminate parental vegetative cells. After ethanol treatment, the spores were centrifuged at 3000 rpm for 5 min, then resuspended in sterile water. To ensure the absence of viable vegetative cells, the spore suspension was incubated in water at 25°C for two weeks. The spores were then centrifuged at 1600 rpm for 3 min and snap-frozen in liquid nitrogen. The vegetative control cells were obtained by adding 10 μL of spores to 20 mL of fresh YES medium and growing at 32°C with shaking at 180 rpm until an OD600 of 0.5-0.7 was reached. The cells were centrifuged at 1600 rpm for 3 min before being snap-frozen. The early stationary-phase control cells were grown on solid YES plates at 32°C for ∼18 hrs, collected in water, centrifuged at 1600 rpm for 3 min, and snap-frozen. Spores were maintained at 25°C in sterile water throughout the experiment. Samples were collected at defined intervals, two weeks, three months, or five months, and stored at –80°C for downstream analyses.

To assess the effect of heat shock on spore viability and determine the optimal temperature for the stress response, spores were subjected to heat stress at 6 temperatures: 25°C (control), 45°C, 50°C, 52°C, 55°C, and 57°C for 1 hour. Following treatment, 100 spores from each condition were plated on YES agar plates. Colony-forming units (CFUs) were counted after incubation, and viability was calculated as the percentage of colonies formed relative to the 25°C control. Based on these results, 45°C was selected as the heat-shock condition for stress-response analysis. Spores were exposed to 45°C for 1 hour, snap-frozen and processed for transcriptomics and proteomics.

To investigate gene expression in offspring cells derived from spores, 50 µL of spore suspensions of different ages or subjected to heat stress (50°C for 30 min) were inoculated into 20 mL of fresh YES medium and incubated at 32°C with shaking at 180 rpm for 14–16 hrs, until reaching an OD_600_ of 0.5–0.7 (mid-exponential phase). Cultures were then harvested by centrifugation at 1600 rpm for 3 min, and cell pellets were immediately snap-frozen in liquid nitrogen for downstream transcriptomic and proteomic analyses.

### Bar-seq mutant screen for spore viability and resilience

Two independent spore pools from the Bioneer library (ver. 5.0; Bioneer, South Korea) were prepared by crossing an auxotroph (*h+*) and a prototroph (*h-*) deletion library. Mating for all deletion mutants with the same deletion mutants was performed using the RoToR HDA robot (Singer Instruments) on MEA medium (Malecki et al. 2016). After 5 days, the plates were incubated at 42°C for 3 days to kill the vegetative cells and isolate pure spores. Colonies were then washed off the plates with water and pooled together and incubated at 25°C. Samples were collected from spores at various timepoints (2 weeks to 6 months) to track mutant frequencies. To investigate spore heat resistance, a sample of 6-month-old pooled spores was also subjected to heat stress at 55°C for 30 min, a condition that killed approximately 60% of the spores (Lethal dose 60: LD60). To avoid DNA contamination from dead spores, spores were germinated in YES liquid medium at 32°C for 2-3 days until stationary phase before collecting cells for DNA extraction.

Spore viability was assessed for 2 independent pooled libraries (Pool A and Pool B) at the various time points (2 weeks, 2 months, 4 months, 5 months, and 6 months) to evaluate their lifespan. At each time point, 100 spores from each pool were separated using a tetrad microdissector microscope (MSM400, Singer Instruments) and plated onto YES agar. Plates were incubated at 32°C, and the resulting colonies were counted to determine the number of viable spores. Viability was calculated as the percentage of colonies formed out of the total spores plated.

Genomic DNA was extracted from the spore samples using the phenol extraction method (Sambrock et al. 1989). Briefly, cells were lysed in screw-cap tubes using a FastPrep-24 Instrument (MP Biomedicals, UK) with 0.5 mm glass beads (Stratech Scientific, UK), 200 μL of lysis buffer, and 200 μL of phenol:chloroform:isoamyl alcohol (25:24:1, ThermoFisher Scientific, USA). The samples were lysed for four cycles of 20 sec at 7m/sec with cooling on ice between runs. The samples were then centrifuged for 5 min at 14,000 rpm, and the aqueous layer was transferred to a new tube. The phenol extraction was repeated twice more, using 200 µL of chloroform:isoamyl alcohol (24:1) and 5 min of centrifugation at 14,000 rpm. DNA was precipitated by adding one volume of isopropanol and 1/10 volume of 3M sodium acetate. The mixture was incubated at –20°C for at least 2 hrs, and centrifuged for 30 min at 14,000 rpm. The DNA pellets were washed with 200 μL of 70% ethanol, centrifuged for 10 min at 14,000 rpm, and air-dried briefly. Finally, the pellet was resuspended in 100 μL of TE buffer containing 3 μL of RNAse (10mg/mL) and incubated for 10 min at room temperature. DNA quantification was performed using a NanoDrop 2000 (Thermo Fisher Scientific, USA).

### Bar-seq library preparation and sequencing

For bar-seq library preparation, 1 µg of total DNA was used as the starting material. UpTag barcodes were amplified using Phusion® High-Fidelity DNA Polymerase (NEB, UK) with custom-designed primers at a final concentration of 100 nM in a total reaction volume of 50 µL. The PCR conditions were as follows: initial denaturation at 98°C for 1 min, followed by 7 cycles of 98°C for 10 sec, 60°C for 30 sec, and 72°C for 30 sec, and a final extension at 72°C for 7 min. PCR products were purified using the MinElute® PCR Purification Kit (Qiagen, Germany) and eluted in 10 µL of dH□O. A second PCR was performed to add Illumina adaptors using Phusion® High-Fidelity DNA Polymerase (NEB, UK). The reaction mix included 12.5 µL of Phusion master mix, 12.5 µL of purified first PCR product, and 0.5 µL each of NEBNext® dual-index Illumina primers (NEB #E7780S) in a final volume of 25 µL. The PCR conditions were: initial denaturation at 98°C for 1 min, followed by 17 cycles of 98°C for 10 sec and 65°C for 1.5 min, and a final extension at 72°C for 7 min. The resulting libraries were size-selected using two consecutive AMPure XP bead cleanups (NEB, UK) at a 0.9x ratio to remove primer dimers and adaptor fragments. The cleaned libraries were quantified using a Qubit dsDNA HS kit and analyzed on an Agilent 2100 Bioanalyzer. Libraries were pooled at a final concentration of 1.1 pM, with 20% PhiX spike-in to increase sequencing diversity. Sequencing was performed using the Illumina NextSeq 500/550 Mid Output kit v2.5 (75 cycles), generating 75-base-pair paired-end reads. The sequencing was carried out at the UCL Cancer Institute sequencing facility.

Barcode identification and differential gene expression analysis: After sequencing, paired-end reads were assembled using PEAR and PCR duplicates were filtered using Seqkit (Zhang et al. 2014; Shen et al. 2016). The UpTag barcodes were extracted using Barcount (https://github.com/Bahler-Lab/barcount), which mapped reads to their respective flanking sequences. The extracted UpTag barcodes were matched to a table containing UpTag sequences and their corresponding gene deletions to generate a total read count for each identified mutant in the sample.

Differential barcode distribution between samples, based on barcode frequency in the re-growth cultures, was performed with edgeR (version 3.40.2) (Robinson et al. 2010). The upper-quartile normalization method was applied to adjust for differences in library sizes and composition between samples. This method calculates normalization factors by dividing each sample’s counts by the 75th percentile of non-zero counts, ensuring robust scaling while minimizing the influence of outlier expression values. Time was treated as a categorical variable, and the pool factor was included to account for differences in barcode abundance across pools. Read counts were modelled using a negative binomial generalized linear model, with likelihood ratio tests used to calculate p-values for barcode abundance differences between time points. To distinguish long– and short-lived mutants, time points at 2 weeks and 6 months were evaluated with a fold change threshold of |log_2_(FC)| >log_2_(3) and a false discovery rate (FDR) threshold of <0.05.

For resilience analysis, we treated heat shock as a factor and included a pool factor to account for barcode abundance variations across different pools. Normalized counts were log-transformed, and specific contrasts were constructed to compare resilience across conditions. Linear models were then fitted to the log-transformed data, with empirical Bayes moderation used for statistical testing. We identified resistant and sensitive mutants using a fold-change threshold of |log_2_(FC)| >log_2_(3) and a false discovery rate (FDR) threshold of <0.05.

### Chronological lifespan assay for *Schizosaccharomyces pombe*

To determine the lifespan of germinants derived from spores of varying ages and under stress conditions, a high-throughput chronological lifespan (CLS) assay was performed using the robotic colony-forming unit (CFU)–based method (Romila et al. 2021). This approach utilizes a lab-developed automated pipeline for CFU quantification.

After spores from three conditions—young (2 weeks), old (5 months), and heat-shocked (50°C for 30 min)—were germinated in YES medium at 32°C for 20–24 hrs with shaking, cultures were diluted to an OD_600_ of 0.002 in fresh 10 mL YES medium and incubated at 32°C with shaking. After 2 days, when cultures reached stationary phase, Day 0 measurements were taken. From each culture, 150 µL was transferred into the first column of a 96-well plate, followed by 1:3 serial dilution in YES medium using an Integra Assist automated multichannel pipette (Integra Biosciences Ltd.). Diluted samples were spotted in quadruplicate onto YES agar PlusPlates using the RoToR HDA robot in the long-pin 96-density format. Plates were incubated at 32°C for 2–4 days until visible colony growth was observed.

Plate images (Epson V700 scanner) were processed using a custom Unix script and analyzed with the DeadOrAlive R package, which automatically grids the images and estimates CFUs from each spot to generate viability over time (Romila et al. 2021). As a proxy for lifespan, the calculated time point at which viability declined to 5% was used as the endpoint metric and was normalized to the average value obtained from progeny derived from non-stressed, 2-week-old spores. All CLS assays were performed in triplicate, and the entire experiment was independently repeated 3 times to ensure reproducibility and statistical robustness.

### Stress spot tests

After spores from three conditions—young (2 weeks), old (5 months old), and heat-shocked (50°C for 30 min)—were germinated in YES medium and incubated at 32°C for 20–24 hrs with shaking, cultures were adjusted to OD_600_ = 0.2, grown until an increase of 0.6 OD_600_ units was reached, and subsequently diluted back to OD□□□ = 0.2 in the indicated media. For each condition, 5 serial 1:3 dilutions were prepared in sterile water. Aliquots (5 µL) from each dilution were spotted onto YES agar plates (control). For heat stress assays, cultures were incubated at 43° or 45°C for 30 min prior to serial 1:3 dilution and spotting. Plates were incubated at 32°C for 2–3 days and imaged. Growth patterns were compared to assess relative tolerance to oxidative and heat stress.

### Killifish husbandry, diapause induction, and embryo collection

*Nothobranchius furzeri* fish (GRZ and MZCS08/122 strains) were housed at the UCL Fish Facility and the Administrative Panel on Laboratory Animal Care. Experimental procedures were approved by the UK Home Office under the Animals (Scientific Procedures) Act 1986 (Licence number: PP9118171). The fish were kept at 28°C with a 14-hr/10-hr light/dark cycle and circulating system water. The daily exchange rate of the system water was 10%, using reverse osmosis water as the source. The pH and conductivity of the system water were automatically monitored and maintained at 7.0 to 7.5 and 3000 μS, respectively. Males and females were housed separately and allowed to breed when needed. Synchronized killifish embryos were collected within a ∼5-hour breeding window in sandboxes. The sand was strained to collect embryos, most of which were at the 1-2 cell stage. The fertilized embryos were washed with Ringer’s solution (Sigma-Aldrich, cat. nr. 96724, Darmstadt, Germany) to remove any debris or contaminants, then stored in Ringer’s solution with 0.01% methylene blue to prevent bacterial and fungal contamination (Hu et al. 2020). The embryos were incubated at 28°C and examined daily to remove dead embryos.

At 4 days post-fertilization, embryos were disinfected with 0.01% sodium hypochlorite (EMPLURA®, 105614) solution prepared in embryo medium for 5 min, followed by three 2-minute washes with fresh media (Podrabsky 1999). This disinfection procedure was repeated three times. After disinfection, embryos were either kept at 28°C to bypass diapause and actively develop or transferred to 20°C to induce diapause. Diapause embryos were collected two weeks post-induction, while actively developing embryos were collected at seven days post-fertilization (d.p.f.), when they are at a similar developmental stage as those in diapause.

Embryos with bulging black eyeballs were transferred to dry media composed of moist coconut fibres and incubated at 28°C for 8 days, until reaching the golden iris stage, when they were ready to hatch. Hatching was induced by incubating embryos in 1 g/L humic acid solution (Sigma-Aldrich, 53680) for two days. The newly hatched fish were considered fry in the first 5 weeks after hatching. Fry was raised in 0.8-litre tanks and fed newly hatched brine shrimps (Brine Shrimp Direct, 454GR). After week 5, fish were considered adults (males show colour around this time, indicative of sexual maturity). Adults were maintained in 2.8 L tanks and received two daily dry feeds consisting of Gemma Micro, Safe Caviar, and Hikari micro pellets, with particle size and proportions adjusted according to developmental stage; adults also received bloodworm (Gamma Slice Bloodworm Mini) and, for breeding fish, additional artemia (cysts, Special Grade 300, ZM), while nursery fish were fed 4 times daily with dry feed and artemia, with feeding reduced to twice daily on weekends. Diapause embryos were maintained at 20°C in Ringer’s solution throughout the experiment. Samples were collected at defined intervals (2 weeks, 3 months, and 4 months) and stored at –80°C. To ensure that Diapause II embryos remained in a stable dormant state, fresh samples were collected two weeks after confirmed entry into diapause. As a control group, actively developing embryos were maintained at 28°C and collected at a comparable developmental stage, specifically, 7 days post-fertilization, shortly after the onset of heartbeat.

To assess the effect of heat shock on embryo viability and determine the optimal temperature for stress-response profiling, both diapause and actively developing embryos were subjected to heat stress at 28°C, 36°C, 38°C, 40°C, and 42°C for 1 hr. Embryos were then returned to their respective maintenance conditions and monitored for survival over the following day. Based on viability results, 40°C was selected as the heat-shock condition for stress-response analysis. Embryos were exposed to 40°C for 1 hr and subsequently collected for RNA-seq and proteomic profiling.

To assess the presence of molecular memory following dormancy, we reactivated embryos from each diapause age group and heat-shock condition by shifting them to 28°C. Ten embryos per group were allowed to resume development for one week, until they reached the pre-hatching stage. These post-dormant embryos were then analyzed to determine whether transcriptomic and proteomic signatures associated with dormancy and stress were retained after reactivation, providing insight into potential molecular memory effects.

For transcriptomic and proteomic analyses of diapause compared to developing embryos, approximately 50 embryos were pooled for RNA-seq and ∼10 embryos per pool for mass spectrometry, with three independent repeats collected at different times. Embryos were dissected in ice-cold PBS using biological-grade tweezers (Electron Microscopy Sciences, 72700-849 D), removing the chorion, enveloping layer, and yolk. Embryo bodies were then snap-frozen in liquid nitrogen and stored at –80°C (Hu et al. 2020).

### RNA extraction and RNA-seq analyses

For RNA extraction, 200 μL 1.0 mm zirconia beads (Cat. No 11079110zx, BioSpec) for diapause and glass 300μm beads (SKU G9143-500G, Sigma) for yeast samples, and 500 μL Qiazole were added to the frozen cell/spore pellets and lysed with a FastPrep® 24 homogeniser at speed 6 m/s for 20 sec 3 times followed by centrifugation at 17,000g for 3 min. The total RNA in the supernatant was purified using the miRNeasy Mini Kit (Qiagen, 217004) according to the manufacturer’s instructions. RNA integrity and concentration were determined using an Agilent 2100 Bioanalyzer and Agilent’s RNA 6000 Pico Kit (Agilent Technologies, no 5067-1513).

RNA-seq libraries were prepared by Genewiz LLC using the NEBNext Ultra II Directional RNA Library Prep Kit for Illumina (NEB, Ipswich, MA). mRNAs were enriched with Oligo(dT) beads, fragmented, and converted to cDNA. The indexed adapter was ligated to cDNA fragments, and limited-cycle PCR was used for amplification. Sequencing was performed on the Illumina NovaSeq 6000 using a 2×150 Paired-End (PE) configuration.

Raw RNA-seq data (FASTQ files) from the killifish diapause or fission yeast RNA-seq experiments were processed and quality-controlled using FastQC v2.4.1. Reads were aligned to the reference genome using Hisat2 v2.2.1 with default parameters (D. Kim et al. 2019). Trimming of 5’ and 3’ bases was performed to remove adaptors. The resulting SAM file was sorted and stored as a BAM file using Samtools v1.9. PCR duplicates were removed using Picard v2.25.1, and read counts for each gene were obtained using featureCounts. Genes with an average count per million (CPM) <1 in more than half of the samples were excluded from further analysis to reduce background noise. Differentially expressed genes (DEGs) were defined as those with an absolute log□ fold change (|log□FC|) >0.58, corresponding to an approximate 1.5-fold change, and a false discovery rate (FDR) <0.05. Differential gene expression analysis was performed using the DESeq2 package (v3.4.1) in R (v4.1.0) (Love et al. 2014). Functional enrichment analysis of DEGs was conducted using the g:Profiler2 package in R (Raudvere et al. 2019). Principal components analysis (PCA) plots were generated using the factoextra R package v1.0.8, and heat maps were created using the gplots R package v3.1.3. For comparison to quiescent cells, the raw data from a previous paper (Marguerat et al. 2012) were used, and the batch effects were removed using combatseq (Zhang et al. 2020).

### Protein sample preparation for proteomics experiments

Killifish embryos were resuspended in 50 μL lysis buffer (final concentration of 5% SDS, 100 mM HEPES and 50 mM DTT). The samples were sonicated using a Bioruptor Plus (Diagenode, Belgium) for 10 cycles (30 sec ON/60 sec OFF) at the high setting, maintaining a temperature of 20°C. The samples were then boiled at 95°C for 10 min. Protein reduction was carried out by alkylation with iodoacetamide (Promega, VB1010) at a final concentration of 15 mM for 30 min at room temperature in the dark. Subsequently, proteins were precipitated by adding 4 volumes of ice-cold 100% acetone to the lysates. The suspensions were centrifuged at 20,000 × g for 30 min at 4°C. The resulting protein pellets were washed three times with 80% acetone, each followed by centrifugation at 20,000 × g for 10 min, and then air-dried. The dried protein pellets were resuspended in digestion buffer (100 mM HEPES, 1 M GuaHCl, pH 8) to a final concentration of 0.5 µg/µL and sonicated for 10 min. LysC enzyme (Promega, VA1170) was then added to the protein solution at a 1:100 ratio and incubated for 4 hrs at 37°C. Afterwards, trypsin (Promega, V5280) was added to the solution, and the mixture was incubated overnight for complete digestion.

To the frozen pellet of spores and vegetative cells, 200□μl of 0.1□M ammonium bicarbonate in 7□M urea was added, along with approximately 100□mg of glass beads per well. The plates were then sealed with cap mats (Spex, no. 2201) and subjected to bead-beating for 5□min at 1,500□r.p.m. using a Spex Geno/Grinder. After lysis, the plates were centrifuged for 1 minute at 4,000□r.p.m., and 20□μl of 55□mM DL-dithiothreitol (final concentration 5□mM) was added with mixing. The samples were incubated for 1 hour at 30□°C. Next, 20□μl of 120□mM iodoacetamide (final concentration 10□mM) was introduced, and the mixture was incubated for 30 min at room temperature in the dark. One millilitre of 100□mM ammonium bicarbonate was subsequently added, and the mixture was centrifuged for 3 min at 4,000□r.p.m. From the resulting supernatant, 230□μl was transferred to trypsin-preloaded plates (9□μl of 0.1□μg□μl–1 trypsin per well). The samples were incubated overnight at 37°C, after which 24□μl of 10% formic acid was added to quench the reaction.

The digestion mixtures were then subjected to solid-phase extraction using C18 96-well plates (96-Well MACROSpin C18, 50–450□μl, The Nest Group, no. SNS SS18VL). Liquid flow through the stationary phase was facilitated by 1-minute centrifugation steps using an Eppendorf Centrifuge 5810□R at varying speeds as described. A liquid handler (Biomek NXP) was used to pipette the solutions, processing four 96-well plates per batch. The plates were conditioned with 200□μl of methanol, then centrifuged at 50g; two washes with 200□μl of 50% acetonitrile (ACN) were performed, with the flow-through discarded after each. Plates were then equilibrated three times with 200□μl of 3% ACN and 0.1% formic acid, centrifuged sequentially at 50g, 80g, and 100g, and discarding the flow-through after each step. Next, 200□μl of the digested samples was loaded onto the plates and centrifuged at 100g, followed by three washes with 200□μl of 3% ACN and 0.1% formic acid, each centrifuged at 100g. After the final wash, the plates were centrifuged again at 180g prior to elution. Peptides were eluted in three steps: twice with 120□μl and once with 130□μl of 50% ACN, each centrifuged at 180g, and collected into a 1.1□ml square-well, V-bottom collection plate. The collected material was then dried using a vacuum concentrator (Eppendorf Concentrator Plus) and redissolved in 40□μl of 3% ACN and 0.1% formic acid, before being transferred to a 96-well plate (700□μl round, Waters, no. 186005837).

### Mass-spectrometry analysis for killifish diapause and developing embryos

Peptides were re-dissolved in Optima™ water/0.1% formic acid. Evotips (Evosep, EV2001 or EV2011) were conditioned according to the manufacturer’s instructions, and the peptide samples and iRT standards (Biognosys AG) were loaded onto conditioned Evotips and queued on an Evosep One liquid chromatography system (Evosep, Denmark) with the pre-defined 60 SPD (samples per day) method and the corresponding Evosep Performance columns (Evosep, EV1109 and EV1137). Mobile phases A and B were 0.1% (v/v) formic acid in water and 0.1% (v/v) formic acid in acetonitrile, respectively. The LC was coupled to a hybrid dual-pressure linear ion trap mass spectrometer (Orbitrap Lumos, Thermo Scientific, San Jose, USA) using an EvoSep EV1072 EasySpray adapter (Evosep, Denmark). Data for all samples was acquired in Data Independent Acquisition mode (DIA). DIA Lumos settings were as follows: Transfer capillary set to 300°C and 2.2 kV applied to the nanospray needle (Evosep, Denmark). MS1 data were acquired in the Orbitrap with a resolution of 120,000 FWHM, max injection time of 20 ms, AGC target of 1×10^6^, in positive ion mode and profile mode, over the mass range 393–907 m/z. DIA segments over this mass range (20 m/z wide/1Da overlap/27 in total) were acquired in the Orbitrap following fragmentation in the HCD cell (32%), with 30k resolution over the mass range 200–2000 m/z, max injection time of 54 ms (dynamic), and AGC target of 1×10^6^.

The data were searched using the Pulsar search engine in Spectronaut (v.14, Biognosys AG). A spectral library was first generated by searching the data against a species-specific in-house database and a database of common contaminants. BGS factory settings (default) were used, except that no fixed modifications were selected. Run-wise imputation (Q-value percentile = 30%) was applied to the dataset.

### Mass-spectrometry analysis for killifish early, late and stressed diapause embryos and post-diapause embryos

Peptides were separated using the Evosep One system (Evosep, Odense, Denmark) equipped with an 8 cm x 150 μm column, packed with 1.5 μm Reprosil-Pur C18 beads (Evosep Endurance, EV-1106, PepSep, Marslev, Denmark). The samples were run with a pre-programmed proprietary Evosep gradient of 21 min (60 samples per day) using water and 0.1% formic acid, and solvent B (acetonitrile) with 0.1% formic acid. The LC was coupled to an Orbitrap Exploris 480 (Thermo Fisher Scientific, Bremen, Germany) using the PepSep Sprayers and Proxeon nanospray source. Peptides were introduced into the mass spectrometer via a PepSep Emitter (outer diameter 360 μm, inner diameter 20 μm), heated to 300°C, with a spray voltage of 2.2 kV applied. The injection capillary temperature was set at 300°C. The radio frequency ion funnel was set to 30%. For data-independent acquisition (DIA), full-scan mass spectrometry (MS) spectra with a mass range of 350–1650 m/z were acquired in profile mode on an Orbitrap at a resolution of 120,000 FWHM. The default charge state was set to 2+. The filling time was set to a maximum of 45 ms, with a limit of 3 x 10^6^ ions. DIA scans were acquired with 35 mass-window segments of varying widths across the MS1 mass range. Higher collisional dissociation fragmentation (stepped normalized collision energy; 25, 27.5, and 30%) was applied, and MS/MS spectra were acquired with a resolution of 15,000 FWHM with a fixed first mass of 200 m/z after accumulation of 1 × 10^6^ ions or after filling time of 37 ms (whichever occurred first). Data were acquired in profile mode. For data acquisition and raw data processing, Xcalibur 4.5 (Thermo) and Tune version 4.0 were used.

For proteomic data processing, DIA raw data were analyzed using the directDIA pipeline in Spectronaut (v.17, Biognosysis, Switzerland). The data were searched against a species-specific in-house database (59,154) and a contaminant SwissProt database (247 entries). The data were probed with the following variable modifications: oxidation (M) and acetyl (protein N-term). A maximum of 2 missed cleavages for trypsin and 5 variable modifications were allowed. The identifications were filtered to satisfy an FDR of 1% at the peptide and protein levels. Relative quantification was performed in Spectronaut for each paired comparison using the replicate samples from each condition. The data (candidate table) and data reports (protein quantities) were then exported, and further data analyses were performed with R using limma workflow (v3.48.3) (Ritchie et al. 2015). For comparison to quiescent cells, the normalized proteome per cell data from a previous study were used (Marguerat et al. 2012).

### Mass-spectrometry analysis for yeast cells

For analyzing the digested proteins, a Waters ACQUITY UPLC M-Class µSystem was used online with a SCIEX ZenoTOF 7600 System mass spectrometer. Prior to analysis, 200 ng of peptides were chromatographically separated at a 5 µL/min flow rate over a 20 min gradient (column: Waters HSS T3, 300µm x 150mm, 1.8µm), heated to 40°C. Mobile phases A and B were 0.1% formic acid in water and 0.1% formic acid in acetonitrile, respectively. For data-independent acquisition, the instrument used a Zeno SWATH MS/MS scheme with 85 variable-size windows and an accumulation time of 11 ms. The ion source parameters were set as follows: ion source gas 1 and gas 2 at 12 psi and 60 psi, respectively; curtain gas 55, CAD gas 7, and source temperature at 200°C; spray voltage at 4500V.

Raw data were processed with DIA-NN (version beta1.8; Demichev et al. 2020) using the default settings except with mass accuracy set to 15 ppm and 15 ppm at the MS2 and MS1 level, respectively, scan window set to 7, MBR (Match-between-runs) enabled, and quantification strategy set as “Robust LC (high Precision)”. A library-free search was performed by first creating an in silico-predicted spectral library from the sequence database: *S. pombe* UniProt fasta (UP000002485, downloaded on 09.03.2022), and then analyzing with this library (Zelezniak et al. 2018). For the annotation, the same database was used, and the output was filtered at 1% FDR on the peptide level.

### Human cancer cell data

We used available RNA-seq data from dormant and active Acute Lymphocytic Leukaemia (ALL) blasts, obtained from children enrolled in clinical multicenter studies (Ebinger et al. 2016). The cells were labeled with BrdU and CFSE and transplanted into NSG mice (NOD/scid, IL2 receptor gamma chain knockout mice). After transplantation, the loss of CFSE in PDX cells was used to distinguish between slowly and rapidly growing subpopulations, referred to as label-retaining cells (LRC) and non-label-retaining cells (non-LRC), respectively. Both LRC and non-LRC cells underwent RNA sequencing. We reanalyzed the raw count matrix using DESeq2, applying a cutoff of log_2_FC >|0.58| and FDR <0.05, with raw data normalized using DESeq (Anders and Huber 2010; Wen 2017).

### Orthology analysis

Orthology relationships between genes in fission yeast and human were obtained from a manually curated PomBase list. Orthology relationships between genes from killifish and human were obtained from a published mapping table (Kelmer Sacramento et al. 2020). In this table, the turquoise killifish proteome was directly mapped to the human UniProt reference proteome using BLASTp, selecting the best matches based on sequence similarity. Additionally, potential hidden orthologs were identified by aligning the turquoise killifish proteome against the spotted gar (*Lepisosteus oculatus*) proteome, with these orthologs subsequently mapped onto the human proteome.

To define the core list of genes involved in dormancy across different model organisms, all genes were mapped to their human orthologs, and all paralogs were included to ensure comprehensive representation. Conserved genes were identified by filtering for protein-coding transcripts that were significantly induced or repressed, with a log_2_FC >|0.58| and a Benjamini-Hochberg-adjusted P-value <0.05 across all models studied. The significance of the overlap was calculated using hypergeometric tests. Overlapping conserved genes were identified by applying the same filters, considering only proteins with orthologs in all three datasets.

### Comparative analysis of process-level expression changes

To evaluate overall process-level expression changes across different biological conditions (diapause, spores, and cancer), we first mapped genes from *N. furzeri* and *S. pombe* to their human orthologs using Ensembl annotations. Gene-level differential expression data (log fold change and adjusted p-values) from RNA-seq and proteomics datasets were then aggregated. Using Ensembl-to-gene symbol mapping and Gene Ontology (GO) Biological Process annotations, genes were assigned to processes, and the mean log_2_FC for each biological process (term_id) was computed per condition. Only processes with sufficient gene representation and variable expression (i.e., ≥2 out of 3 conditions showing absolute log_2_FC >0.3) were retained for analysis. We calculated pairwise Pearson correlations of these process-level log_2_FC values across conditions, providing a comparative view of expression trends at both the transcriptomic and proteomic levels.

## Data availability

The RNA-seq data generated in this study have been deposited in the European Nucleotide Archive (ENA) under accession number PRJEB80774. Proteomics datasets generated from yeast spores, and killifish MZCS early, late and stressed diapause embryos and post-diapause embryos were deposited in the PRIDE repository under accession numbers PXD077825 and PXD077884, respectively, and those generated from killifish GRZ diapause vs developing embryos were deposited to the MassIVE repository under accession number MSV000101686.

## Acknowledgments

We thank Joanna Kirkpatrick and Mark Skehel of the Proteomics Facility at the Francis Crick Institute and Anja Freiwald and Kathrin Textoris-Taube of the Core Facility High Throughput Mass Spectrometry, Charité – Universitätsmedizin Berlin, and Therese Dau and Emilio Cirri from the FLI Core Facility Proteomics, for their valuable support with proteomics sample preparation, data acquisition, and analysis. We thank Heather Callaway, Paul Barwood, Joe Warmsley, and other staff of the UCL Fish Facility for their valuable support with killifish husbandry and maintenance, and Giovanni Stefani for the efforts to get our killifish colony initially established. We thank Ben Heineke and Jennifer Stiens for their critical reading of the manuscript. We used Grammarly to refine some of the writing and q.e.d Science to highlight potential limitations. This research was supported by a Wellcome Discovery Award (302608/Z/23/Z), a Leverhulme Trust Research Project Grant (RPG-2022-067), and a Biotechnology and Biological Sciences Research Council grant (BB/R009597/1) to J.B., a Scholarship from the Newton-Mosharafa Fund to S.H., and a studentship from the Turkish Ministry of National Education to B.K.

## References

1. Anders, Simon, and Wolfgang Huber. 2010. ‘Differential Expression Analysis for Sequence Count Data’. Genome Biology 11 (10): 1–12. 10.1186/GB-2010-11-10-R106/COMMENTS.

2. Baumgart, Mario, Marco Groth, Steffen Priebe, et al. 2014. ‘RNA-Seq of the Aging Brain in the Short-Lived Fish N. Furzeri – Conserved Pathways and Novel Genes Associated with Neurogenesis’. Aging Cell 13 (6): 965–74. 10.1111/acel.12257.

3. Billmyre, R. Blake, Michael T. Eickbush, Caroline J. Craig, et al. 2022. ‘Genome-Wide Quantification of Contributions to Sexual Fitness Identifies Genes Required for Spore Viability and Health in Fission Yeast’. PLOS Genetics 18 (10): e1010462. 10.1371/journal.pgen.1010462.

4. Braun, Sigurd, Cornelia Kilchert, Aydan Bulut-Karslioglu, et al. 2026. ‘Nuclear Dynamics in Quiescent Cells: Conserved Mechanisms from Yeasts to Mammals’. Biomolecules 16 (2): 203. 10.3390/biom16020203.

5. Brunet, Anne. 2020. ‘Old and New Models for the Study of Human Ageing’. Nature Reviews. Molecular Cell Biology 21 (9): 491–93. 10.1038/S41580-020-0266-4.

6. Cellerino, Alessandro, Dario R. Valenzano, and Martin Reichard. 2016. ‘From the Bush to the Bench: The Annual Nothobranchius Fishes as a New Model System in Biology’. Biological Reviews of the Cambridge Philosophical Society 91 (2): 511–33. 10.1111/BRV.12183.

7. Coller, Hilary A. 2011. ‘The Essence of Quiescence’. Science 334 (6059): 1074–75. 10.1126/science.1216242.

8. Demichev, Vadim, Christoph B. Messner, Spyros I. Vernardis, Kathryn S. Lilley, and Markus Ralser. 2020. ‘DIA-NN: Neural Networks and Interference Correction Enable Deep Proteome Coverage in High Throughput’. Nature Methods 17 (1): 41–44. 10.1038/s41592-019-0638-x.

9. Di Fraia, Domenico, Antonio Marino, Jae Ho Lee, et al. 2025. ‘Altered Translation Elongation Contributes to Key Hallmarks of Aging in the Killifish Brain’. Science 389 (6759): eadk3079. 10.1126/science.adk3079.

10. Dick, Fiona, Ole Bjørn Tysnes, Guido W. Alves, Gonzalo S. Nido, and Charalampos Tzoulis. 2023. ‘Altered Transcriptome-Proteome Coupling Indicates Aberrant Proteostasis in Parkinson’s Disease’. iScience 26 (2): 105925. 10.1016/J.ISCI.2023.105925.

11. Dolfi, Luca, Roberto Ripa, Adam Antebi, Dario Riccardo Valenzano, and Alessandro Cellerino. 2019. ‘Cell Cycle Dynamics during Diapause Entry and Exit in an Annual Killifish Revealed by FUCCI Technology’. EvoDevo 10 (1): 1–22. 10.1186/S13227-019-0142-5/FIGURES/14.

12. Ebinger, Sarah, Erbey Ziya Özdemir, Christoph Ziegenhain, et al. 2016. ‘Characterization of Rare, Dormant, and Therapy-Resistant Cells in Acute Lymphoblastic Leukemia’. Cancer Cell 30 (6): 849–62. 10.1016/J.CCELL.2016.11.002.

13. Elkin, Adam M., Sarah Robbins, Claudia S. Barros, and Torsten Bossing. 2025. ‘The Critical Balance Between Quiescence and Reactivation of Neural Stem Cells’. Biomolecules 15 (5): 672. 10.3390/biom15050672.

14. Errington, Jeff. 2003. ‘Regulation of Endospore Formation in Bacillus Subtilis’. Nature Reviews. Microbiology 1 (2): 117–26. 10.1038/NRMICRO750.

15. Fielenbach, Nicole, and Adam Antebi. 2008. ‘C. Elegans Dauer Formation and the Molecular Basis of Plasticity’. Genes & Development 22 (16): 2149–65. 10.1101/GAD.1701508.

16. Furness, Andrew I., David N. Reznick, Mark S. Springer, and Robert W. Meredith. 2015. ‘Convergent Evolution of Alternative Developmental Trajectories Associated with Diapause in African and South American Killifish’. Proceedings of the Royal Society B: Biological Sciences 282 (1802): 20142189. 10.1098/rspb.2014.2189.

17. Gemin, Olivier, Maciej Gluc, Higor Rosa, et al. 2024. ‘Ribosomes Hibernate on Mitochondria during Cellular Stress’. Nature Communications 15 (1): 8666. 10.1038/s41467-024-52911-4.

18. Gottlieb, S., and G. Ruvkun. 1994. ‘Daf-2, Daf-16 and Daf-23: Genetically Interacting Genes Controlling Dauer Formation in Caenorhabditis Elegans’. Genetics 137 (1): 107–20. 10.1093/GENETICS/137.1.107.

19. Han, Tian X., Xing Ya Xu, Mei Jun Zhang, Xu Peng, and Li Lin Du. 2010. ‘Global Fitness Profiling of Fission Yeast Deletion Strains by Barcode Sequencing’. Genome Biology 11 (6): 1–13. 10.1186/GB-2010-11-6-R60/FIGURES/6.

20. Hand, Steven C., David L. Denlinger, Jason E. Podrabsky, and Richard Roy. 2016. ‘Mechanisms of Animal Diapause: Recent Developments from Nematodes, Crustaceans, Insects, and Fish’. American Journal of Physiology. Regulatory, Integrative and Comparative Physiology 310 (11): R1193–211. 10.1152/AJPREGU.00250.2015.

21. Harel, Itamar. 2022. ‘The Turquoise Killifish’. Nature Methods 19: pages 1150–1151.

22. Hu, Chi Kuo, Wei Wang, Julie Brind’Amour, et al. 2020. ‘Vertebrate Diapause Preserves Organisms Long Term through Polycomb Complex Members’. Science (New York, N.Y.) 367 (6480): 870. 10.1126/SCIENCE.AAW2601.

23. Janssens, Georges E., Anne C. Meinema, Javier González, et al. 2015. ‘Protein Biogenesis Machinery Is a Driver of Replicative Aging in Yeast’. eLife 4 (September): e08527. 10.7554/eLife.08527.

24. Jiang, Qi, Michelle Ertel, Austin Arrigo, et al. 2025. ‘Role of the NuRD Complex and Altered Proteostasis in Cancer Cell Quiescence’. Preprint, bioRxiv, February 14. 10.1101/2025.02.10.637435.

25. Kelmer Sacramento, Erika, Joanna M. Kirkpatrick, Mariateresa Mazzetto, et al. 2020. ‘Reduced Proteasome Activity in the Aging Brain Results in Ribosome Stoichiometry Loss and Aggregation’. Molecular Systems Biology 16 (6). 10.15252/msb.20209596.

26. Kim, Daehwan, Joseph M. Paggi, Chanhee Park, Christopher Bennett, and Steven L. Salzberg. 2019. ‘Graph-Based Genome Alignment and Genotyping with HISAT2 and HISAT-Genotype’. Nature Biotechnology 37 (8): 907–15. 10.1038/s41587-019-0201-4.

27. Kim, Dong-Uk, Jacqueline Hayles, Dongsup Kim, et al. 2010. ‘Analysis of a Genome-Wide Set of Gene Deletions in the Fission Yeast Schizosaccharomyces Pombe’. Nature Biotechnology 28 (6): 617–23. 10.1038/nbt.1628.

28. Li, Nheng, and Hans Clevers. 2010. ‘Coexistence of Quiescent and Active Adult Stem Cells in Mammals’. Science (New York, N.Y.) 327 (5965): 542–45. 10.1126/SCIENCE.1180794.

29. López-Maury, Luis, Samuel Marguerat, and Jürg Bähler. 2008. ‘Tuning Gene Expression to Changing Environments: From Rapid Responses to Evolutionary Adaptation’. Nature Reviews Genetics 2008 9:8 9 (8): 583–93. 10.1038/nrg2398.

30. López-Otín, Carlos, Maria A. Blasco, Linda Partridge, Manuel Serrano, and Guido Kroemer. 2023. ‘Hallmarks of Aging: An Expanding Universe’. Cell 186 (2): 243–78. 10.1016/j.cell.2022.11.001.

31. Love, Michael I., Wolfgang Huber, and Simon Anders. 2014. ‘Moderated Estimation of Fold Change and Dispersion for RNA-Seq Data with DESeq2’. Genome Biology 15 (12): 550. 10.1186/s13059-014-0550-8.

32. MacRae, Thomas H. 2010. ‘Gene Expression, Metabolic Regulation and Stress Tolerance during Diapause’. Cellular and Molecular Life Sciences: CMLS 67 (14): 2405–24. 10.1007/s00018-010-0311-0.

33. Maire, Théo, Tim Allertz, Max A. Betjes, and Hyun Youk. 2020. ‘Dormancy□to□death Transition in Yeast Spores Occurs Due to Gradual Loss of Gene□expressing Ability’. Molecular Systems Biology 16 (11): 9245. 10.15252/MSB.20199245.

34. Malecki, Michal, Danny A. Bitton, Maria Rodríguez-López, et al. 2016. ‘Functional and Regulatory Profiling of Energy Metabolism in Fission Yeast’. Genome Biology 17 (1): 1–18. 10.1186/S13059-016-1101-2/FIGURES/8.

35. Marcelo, Adriana, Rebekah Koppenol, Luís Pereira de Almeida, Carlos A. Matos, and Clévio Nóbrega. 2021. ‘Stress Granules, RNA-Binding Proteins and Polyglutamine Diseases: Too Much Aggregation?’ Cell Death & Disease 12 (6): 592. 10.1038/s41419-021-03873-8.

36. Marescal, Océane, and Iain M. Cheeseman. 2020. ‘Cellular Mechanisms and Regulation of Quiescence’. Developmental Cell 55 (3): 259–71. 10.1016/j.devcel.2020.09.029.

37. Marguerat, Samuel, Alexander Schmidt, Sandra Codlin, Wei Chen, Ruedi Aebersold, and Jürg Bähler. 2012. ‘Quantitative Analysis of Fission Yeast Transcriptomes and Proteomes in Proliferating and Quiescent Cells’. Cell 151 (3): 671–83. 10.1016/J.CELL.2012.09.019.

38. Mata, Juan, Rachel Lyne, Gavin Burns, and Jürg Bähler. 2002. ‘The Transcriptional Program of Meiosis and Sporulation in Fission Yeast’. Nature Genetics 32 (1): 143–47. 10.1038/NG951.

39. Mata, Juan, Anna Wilbrey, and Jürg Bähler. 2007. ‘Transcriptional Regulatory Network for Sexual Differentiation in Fission Yeast’. Genome Biology 8 (10): R217. 10.1186/gb-2007-8-10-r217.

40. Mead, R. A. 1981. ‘Delayed Implantation in Mustelids, with Special Emphasis on the Spotted Skunk.’ Journal of Reproduction and Fertility. Supplement 29 (January): 11–24.

41. Morree, Antoine de, and Thomas A. Rando. 2023. ‘Regulation of Adult Stem Cell Quiescence and Its Functions in the Maintenance of Tissue Integrity’. Nature Reviews. Molecular Cell Biology 24 (5): 334–54. 10.1038/s41580-022-00568-6.

42. Mukaiyama, Hiroyuki, Shiro Kajiwara, Akira Hosomi, et al. 2009. ‘Autophagy-Deficient Schizosaccharomyces Pombe Mutants Undergo Partial Sporulation during Nitrogen Starvation’. *Microbiology* (Reading, England) 155 (Pt 12): 3816–26. 10.1099/mic.0.034389-0.

43. Murley, Andrew, Ann Catherine Popovici, Xiwen Sophie Hu, et al. 2025. ‘Quiescent Cell Re-Entry Is Limited by Macroautophagy-Induced Lysosomal Damage’. Cell 188 (10): 2670–2686.e14. 10.1016/j.cell.2025.03.009.

44. Nonninger, Tim J., Jennifer Mak, Birgit Gerisch, et al. 2025. ‘A TFEB–TGFβ Axis Systemically Regulates Diapause, Stem Cell Resilience and Protects against a Senescence-like State’. Nature Aging 5 (7): 1340–57. 10.1038/s43587-025-00911-4.

45. Nuckolls, Nicole L., Michael T. Eickbush, Jeffrey J. Lange, Christopher J. Wood, Stephanie H. Nowotarski, and Sarah E. Zanders. 2025. ‘Impacts of Stress and Aging on Spore Health in Schizosaccharomyces Pombe’. Microbiology Spectrum 13 (12): e0171025. 10.1128/spectrum.01710-25.

46. Oh, Juhyun, Yang David Lee, and Amy J. Wagers. 2014. ‘Stem Cell Aging: Mechanisms, Regulators and Therapeutic Opportunities’. Nature Medicine 20 (8): 870–80. 10.1038/NM.3651.

47. Ohtsuka, Hokuto, Kazuki Imada, Takafumi Shimasaki, and Hirofumi Aiba. 2022. ‘Sporulation: A Response to Starvation in the Fission Yeast Schizosaccharomyces Pombe’. MicrobiologyOpen 11 (3): e1303. 10.1002/MBO3.1303.

48. Phan, Tri Giang, and Peter I. Croucher. 2020. ‘The Dormant Cancer Cell Life Cycle’. Nature Reviews Cancer 2020 20:7 20 (7): 398–411. 10.1038/s41568-020-0263-0.

49. Platzer, Matthias, and Christoph Englert. 2016. ‘Nothobranchius Furzeri: A Model for Aging Research and More’. Trends in GeneticslJ: TIG 32 (9): 543–52. 10.1016/J.TIG.2016.06.006.

50. Podrabsky, Jason E. 1999. ‘Husbandry of the Annual Killifish Austrofundulus Limnaeus with Special Emphasis on the Collection and Rearing of Embryos’. Environmental Biology of Fishes 54 (4): 421–31. 10.1023/A:1007598320759/METRICS.

51. Poeschla, Michael, and Dario R. Valenzano. 2020. ‘The Turquoise Killifish: A Genetically Tractable Model for the Study of Aging’. Journal of Experimental Biology 223 (February). 10.1242/JEB.209296.

52. Polačik, M., R. Blažek, R. Řežucha, M. Vrtílek, E. Terzibasi Tozzini, and M. Reichard. 2014. ‘Alternative Intrapopulation Life□history Strategies and Their Trade□offs in an A Frican Annual Fish’. Journal of Evolutionary Biology 27 (5): 854–65. 10.1111/jeb.12359.

53. Ptak, Grazyna E., Emanuela Tacconi, Marta Czernik, Paola Toschi, Jacek A. Modlinski, and Pasqualino Loi. 2012. ‘Embryonic Diapause Is Conserved across Mammals’. PLoS ONE 7 (3): 33027. 10.1371/JOURNAL.PONE.0033027.

54. Raudvere, Uku, Liis Kolberg, Ivan Kuzmin, et al. 2019. ‘G:Profiler: A Web Server for Functional Enrichment Analysis and Conversions of Gene Lists (2019 Update)’. Nucleic Acids Research 47 (W1): W191–98. 10.1093/nar/gkz369.

55. Reichwald, Kathrin, Andreas Petzold, Philipp Koch, et al. 2015. ‘Insights into Sex Chromosome Evolution and Aging from the Genome of a Short-Lived Fish’. Cell 163 (6): 1527–38. 10.1016/J.CELL.2015.10.071.

56. Renfree, Marilyn B., and Jane C. Fenelon. 2017. ‘The Enigma of Embryonic Diapause’. *Development (Cambridge*, England) 144 (18): 3199–210. 10.1242/dev.148213.

57. Ritchie, Matthew E., Belinda Phipson, Di Wu, et al. 2015. ‘Limma Powers Differential Expression Analyses for RNA-Sequencing and Microarray Studies’. Nucleic Acids Research 43 (7). 10.1093/nar/gkv007.

58. Rittershaus, Emily S. C., Seung Hun Baek, and Christopher M. Sassetti. 2013. ‘The Normalcy of Dormancy: Common Themes in Microbial Quiescence’. Cell Host & Microbe 13 (6): 643–51. 10.1016/J.CHOM.2013.05.012.

59. Robinson, Mark D., Davis J. McCarthy, and Gordon K. Smyth. 2010. ‘edgeR: A Bioconductor Package for Differential Expression Analysis of Digital Gene Expression Data’. Bioinformatics 26 (1): 139–40. 10.1093/bioinformatics/btp616.

60. Roche, Benjamin, Benoit Arcangioli, and Robert Martienssen. 2017. ‘Transcriptional Reprogramming in Cellular Quiescence’. RNA Biology 14 (7): 843. 10.1080/15476286.2017.1327510.

61. Romila, Catalina A., St John Townsend, Michal Malecki, et al. 2021. ‘Barcode Sequencing and a High-Throughput Assay for Chronological Lifespan Uncover Ageing-Associated Genes in Fission Yeast’. Microbial Cell 8 (7): 146. 10.15698/MIC2021.07.754.

62. Rossi, Alice, Antoine Coum, Manon Madelenat, et al. 2026. ‘Cellular Quiescence Uncouples the Proteome from the Transcriptome in Neural Stem Cells’. The EMBO Journal 45 (5): 1697–727. 10.1038/s44318-026-00693-4.

63. Sakai, Keiichiro, Yohei Kondo, Kazuhiro Aoki, and Yuhei Goto. 2025. ‘Molecular and Biophysical Perspectives on Dormancy Breaking: Lessons from Yeast Spore’. Biomolecules 15 (5): 701. 10.3390/biom15050701.

64. Segev, Einat, Yoav Smith, and Sigal Ben-Yehuda. 2012. ‘RNA Dynamics in Aging Bacterial Spores’. Cell 148 (1–2): 139–49. 10.1016/j.cell.2011.11.059.

65. Shetty, Sunil, Jon Hofstetter, Stefania Battaglioni, Danilo Ritz, and Michael N. Hall. 2023. ‘TORC1 Phosphorylates and Inhibits the Ribosome Preservation Factor Stm1 to Activate Dormant Ribosomes’. The EMBO Journal 42 (5): e112344. 10.15252/embj.2022112344.

66. Sideri, Theodora, Charalampos Rallis, Danny A. Bitton, et al. 2015. ‘Parallel Profiling of Fission Yeast Deletion Mutants for Proliferation and for Lifespan During Long-Term Quiescence’. G3 Genes|Genomes|Genetics 5 (1): 145–55. 10.1534/g3.114.014415.

67. Singh, Param Priya, G. Adam Reeves, Kévin Contrepois, et al. 2024. ‘Evolution of Diapause in the African Turquoise Killifish by Remodeling the Ancient Gene Regulatory Landscape’. Cell 187 (13): 3338–3356.e30. 10.1016/j.cell.2024.04.048.

68. Sun, Ling-Ling, Ming Li, Fang Suo, et al. 2013. ‘Global Analysis of Fission Yeast Mating Genes Reveals New Autophagy Factors’. PLoS Genetics 9 (8): e1003715. 10.1371/journal.pgen.1003715.

69. Takemon, Yuka, Joel M. Chick, Isabela Gerdes Gyuricza, et al. 2021. ‘Proteomic and Transcriptomic Profiling Reveal Different Aspects of Aging in the Kidney’. eLife 10 (March): e62585. 10.7554/eLife.62585.

70. Thompson, Andrew W., and Guillermo Ortí. 2016. ‘Annual Killifish Transcriptomics and Candidate Genes for Metazoan Diapause’. Molecular Biology and Evolution 33 (9): 2391–95. 10.1093/molbev/msw110.

71. Thompson, Dawn, Aviv Regev, and Sushmita Roy. 2015. ‘Comparative Analysis of Gene Regulatory Networks: From Network Reconstruction to Evolution’. Annual Review of Cell and Developmental Biology 31 (Volume 31, 2015): 399–428. 10.1146/annurev-cellbio-100913-012908.

72. Tsuyuzaki, Hayato, Masahito Hosokawa, Koji Arikawa, et al. 2020. ‘Time-Lapse Single-Cell Transcriptomics Reveals Modulation of Histone H3 for Dormancy Breaking in Fission Yeast’. Nature Communications 11 (1): 1265. 10.1038/s41467-020-15060-y.

73. Tsuyuzaki, Hayato, Ryosuke Ujiie, and Masamitsu Sato. 2021. ‘Wake-up Alarm: Virtual Time-Lapse Gene Expression Landscape Illuminates Mechanisms Underlying Dormancy Breaking of Germinating Spores’. Current Genetics 67 (4): 519–34. 10.1007/S00294-021-01177-0.

74. Tyshkovskiy, Alexander, Siming Ma, Anastasia V. Shindyapina, et al. 2023. ‘Distinct Longevity Mechanisms across and within Species and Their Association with Aging’. Cell 186 (13): 2929–2949.e20. 10.1016/j.cell.2023.05.002.

75. Valcourt, James R., Johanna M. S. Lemons, Erin M. Haley, Mina Kojima, Olukunle O. Demuren, and Hilary A. Coller. 2012. ‘Staying Alive: Metabolic Adaptations to Quiescence’. *Cell Cycle (Georgetown*, Tex.) 11 (9): 1680–96. 10.4161/CC.19879.

76. Van Dyke, Natalya, Ekkawit Chanchorn, and Michael W. Van Dyke. 2013. ‘The Saccharomyces Cerevisiae Protein Stm1p Facilitates Ribosome Preservation during Quiescence’. Biochemical and Biophysical Research Communications 430 (2): 745–50. 10.1016/j.bbrc.2012.11.078.

77. Virgilio, Claudio de. 2012. ‘The Essence of Yeast Quiescence’. FEMS Microbiology Reviews 36 (2): 306–39. 10.1111/J.1574-6976.2011.00287.X.

78. Wen, Guangzheng. 2017. ‘A Simple Process of RNA-Sequence Analyses by Hisat2, Htseq and DESeq2’. Proceedings of the 2017 International Conference on Biomedical Engineering and Bioinformatics, September 14, 11–15. 10.1145/3143344.3143354.

79. Wiecek, Anna J., Stephen J. Cutty, Daniel Kornai, et al. 2023. ‘Genomic Hallmarks and Therapeutic Implications of G0 Cell Cycle Arrest in Cancer’. Genome Biology 24 (1): 128. 10.1186/s13059-023-02963-4.

80. Wood, Valerie, Antonia Lock, Midori A. Harris, Kim Rutherford, Jürg Bähler, and Stephen G. Oliver. 2019. ‘Hidden in Plain Sight: What Remains to Be Discovered in the Eukaryotic Proteome?’ Open Biology 9 (2). 10.1098/rsob.180241.

81. Wourms, John P. 1972. ‘The Developmental Biology of Annual Fishes. III. Pre-Embryonic and Embryonic Diapause of Variable Duration in the Eggs of Annual Fishes’. Journal of Experimental Zoology 182 (3): 389–414. 10.1002/jez.1401820310.

82. Yanagida, Mitsuhiro. 2009. ‘Cellular Quiescence: Are Controlling Genes Conserved?’ Trends in Cell Biology 19(12): 705–15. 10.1016/J.TCB.2009.09.006.

83. Yeger-Lotem, Esti, Laura Riva, Linhui Julie Su, et al. 2009. ‘Bridging High-Throughput Genetic and Transcriptional Data Reveals Cellular Responses to Alpha-Synuclein Toxicity’. Nature Genetics 41 (3): 316–23. 10.1038/ng.337.

84. Zelezniak, Aleksej, Jakob Vowinckel, Floriana Capuano, et al. 2018. ‘Machine Learning Predicts the Yeast Metabolome from the Quantitative Proteome of Kinase Knockouts’. Cell Systems 7 (3): 269–283.e6. 10.1016/j.cels.2018.08.001.

85. Zhang, Yuqing, Giovanni Parmigiani, and W. Evan Johnson. 2020. ‘ComBat-Seq: Batch Effect Adjustment for RNA-Seq Count Data’. NAR Genomics and Bioinformatics 2 (3). 10.1093/nargab/lqaa078.

